# Revolutionizing GPCR-Ligand Predictions: DeepGPCR with experimental Validation for High-Precision Drug Discovery

**DOI:** 10.1101/2024.02.25.581988

**Authors:** Haiping Zhang, Hongjie Fan, Jixia Wang, Tao Hou, Konda Mani Saravanan, Wei Xia, Hei Wun Kan, Junxin Li, John Z.H. Zhang, Xinmiao Liang, Yang Chen

**Affiliations:** Faculty of Synthetic Biology and Institute of Synthetic Biology, Shenzhen Institute of Advanced Technology, Chinese Academy of Sciences, Shenzhen 518055, China; Ganjiang Chinese Medicine Innovation Center, Nanchang 330000, China; CAS Key Laboratory of Separation Science for Analytical Chemistry, Dalian Institute of Chemical Physics, Chinese Academy of Sciences, Dalian 116023, China; Department of Biotechnology, Bharath Institute of Higher Education and Research, Chennai 600073, Tamil Nadu, India; Shenzhen Laboratory of Human Antibody Engineering, Institute of Biomedicine and Biotechnology, Shenzhen Institutes of Advanced Technology, Chinese Academy of Sciences, Shenzhen 518055, China

**Keywords:** GPCR, GLP-1R, GPR35, Graph Convolutional Network, Drug screening

## Abstract

G-protein coupled receptors (GPCRs), crucial in various diseases, are targeted of over 40% of approved drugs. However, the reliable acquisition of experimental GPCRs structures is hindered by their lipid-embedded conformations. Traditional protein-ligand interaction models falter in GPCR-drug interactions, caused by limited and low-quality structures. Generalized models, trained on soluble protein-ligand pairs, are also inadequate. To address these issues, we developed two models, DeepGPCR_BC for binary classification and DeepGPCR_RG for affinity prediction. These models use non-structural GPCR-ligand interaction data, leveraging graph convolutional networks (GCN) and mol2vec techniques to represent binding pockets and ligands as graphs. This approach significantly speeds up predictions while preserving critical physical-chemical and spatial information. In independent tests, DeepGPCR_BC surpassed Autodock Vina and Schrödinger Dock with an AUC of 0.72, accuracy of 0.68, and TPR of 0.73, whereas DeepGPCR_RG demonstrated a Pearson correlation of 0.39 and RMSE of 1.34. We applied these models to screen drug candidates for GPR35 (Q9HC97), yielding promising results with 3 (F545-1970, K297-0698, S948-0241) out of 8 candidates. Furthermore, we also successfully obtained 6 active inhibitors for GLP-1R. Our GPCR-specific models pave the way for efficient and accurate large-scale virtual screening, potentially revolutionizing drug discovery in the GPCR field.

## Introduction

G protein-coupled receptors (GPCRs) are a crucial family of membrane proteins that play a critical role in signal transduction, regulating numerous physiological processes in humans, such as neurotransmission, secretion, cellular differentiation, growth, inflammation, and more ^1–3^. Many diseases are associated with GPCRs, and approximately one-third of approved drugs are designed to interact with GPCRs ^4^. However, many diseases associated with GPCRs still lack approved drugs that can effectively modulate them, thus underscoring the enormous potential of GPCRs as novel targets for disease curing. Unfortunately, the limited availability of GPCR-ligand structures (below 500) ^5^ and the difficulty in obtaining reliable GPCR structures because of their embedding in the lipid membrane pose significant challenges for developing structure-based GPCR-ligand prediction models. It is worth noting that the unique properties of GPCRs compared to soluble proteins in protein-ligand binding further complicate the modeling process.

In recent years, deep learning has emerged as a powerful tool for predicting protein-ligand interaction, thanks to the availability of large protein-ligand datasets like PDBbind ^6^ and BindingDB ^7^. Numerous generalized protein-ligand prediction models have been developed, including structure complex-based models like DeepBindBC ^8^, 3D fusion model ^9^, PointNet ^10^, PointTransformer ^10^, and pafnucy ^11^, as well as non-structure complex based models like DFCNN ^12^, DeepLPI ^13^, DeepDTAF ^14^, CAPLA ^15^ and GraphDTA ^16^. However, GPCR proteins are embedded in the lipid membrane, meaning their physical-chemical environment differs from other soluble proteins ^17^. As a result, the characteristics of GPCR-ligand interactions are distinctly different from those of other protein-ligand interactions, and in some cases, the interaction rules are opposite. Currently available generalized protein-ligand models have poor performance on GPCR-ligand prediction, as highlighted in recent research ^12^. For example, the DFCNN model’s virtual screening over GPCR is inferior to other types of proteins and sometimes even demonstrates worse performance than random predictions ^12^. Hence, there is a necessity to develop a GPCR-ligand-specific model that exclusively trains on the GPCR-ligand dataset and implicitly accounts for the lipid effect. Directly training such a model with the limited availability of GPCR-ligand structures is not feasible in the current scenario. Moreover, generalized models prove to be ineffective for GPCR due to the distinctive binding properties of GPCR-ligand interactions.

Therefore, developing a model that can accurately predict GPCR-ligand interactions is highly attractive and necessary. However, the available 3D structures for GPCR-ligand complexes are limited (<500). At the same time, most of the current protein-ligand predictions are based on learning experimental obtained 3D protein-ligand complexes, such as pafnucy ^11^, DeepBindBC ^8^, and DeepBindRG ^18^. Even worse, the quality of GPCR-ligand is poor because of its difficulty in crystallizing; its native state is in lipid membranes with flexible conformation. Some protein-ligand prediction models do not rely on protein-ligand 3D complexes, such as DFCNN ^19^, but DFCNN lacks spatial information about the binding pocket, which may be critical for those GPCR-ligand binding. Since the GPCR-ligand interaction pattern differs from other types of proteins, it would be impractical to use transfer learning for the GPCR-ligand interaction from other types of protein-ligand 3D datasets. Conversely, GPCR-ligand binding information is abundant; for instance, the GLASS database has collected more than 100,000 protein-ligand pairs with affinity information. Considering the above, developing a non-structure based GPCR-ligand interaction model would be an optimal option. Here, we focused on training GPCR-ligand interaction models by implementing the protein pocket and ligand information with the graph representation. By adopting this methodology, the model achieves independence from the 3D protein-ligand complexes.

In this study, we gathered GPCR-compound pairs from the GLASS database ^20^, and focusing on the extracted GPCR structures and their known pockets. Finally, we employed these GPCR-ligand pairs to train and validate our models and obtained DeepGPCR_BC and DeepGPCR_RG for binary classification and affinity prediction. Additionally, we used a separate test set of eight GPCRs with known active compounds, using modeled structures and predicted pockets. Our two models demonstrated high accuracy, indicating their ability to screen potential drugs for GPCRs even when experimental structures are unavailable. In performance metrics, both DeepGPCR_BC and DeepGPCR_RG shown clear advantages over Schrödinger and vina docking methods in terms of performance. We integrated our computational models with docking techniques to identify possible active agents targeting Q9HC97 (GPR35) and GLP-1R. Two distinct strategies were employed: Strategy A involved the use of DeepGCPR_BC, DeepGPCR_RG, and Schrödinger docking, while Strategy B utilized DeepGPCR_RG in conjunction with Schrödinger docking. From the eight candidates chosen through Strategy A for experimental validation, three demonstrated activities against GPR35 (compounds F545-1970, K297-0698, and S948-0241). For GLP-1R, Strategy A yielded five active compounds out of 12 candidates tested, and Strategy B produced one active compound from three tested. Additionally, molecular dynamics (MD) simulations were conducted to investigate the binding dynamics and atomistic interactions of the three active GPR35 compounds. The findings underscore the potential of our method GPCR-targeted drug screening, particularly when integrated with docking and MD simulation techniques.

## Methods

### Data collection

The GPCR-ligand pairs for this study were sourced from the GLASS database ^20^, with ligand molecules in SMILES format converted to 3DSDF format using the RDKit tool ^21^. For DeepGPCR_BC and DeepGPCR_RG, we have different treatments as following:

#### DeepGPCR_BC

GPCR-ligand pairs with binding affinities characterized by IC50, Ki, or Kd values smaller than 4 nM were considered positive data, indicating strong binding affinity. In contrast, pairs with IC50, Ki, or Kd values larger than 4000 nM were considered negative data, indicating weak or non-binding. The corresponding PDB structure for each GPCR uniprot ID was retrieved by ID Mapping^22^ and downloaded from the PDB database^23^. The representative PDB structure with a known ligand was selected for each GPCR. Residues within 0.6 nm of the known ligand were extracted as the protein pocket.

#### DeepGPCR_RG

Binding affinities of GPCR-ligand pairs are used as training labels. The corresponding structures for each GPCR uniport ID were retrieved from the Alphafold database^24^. The predicted ligands were obtained by the COFACTOR^25^, with selection criteria favoring those closest to the N terminus of the GPCR. The N-terminus of a GPCR, typically extracellular, often contains the native ligand binding region. This region is responsible for recognizing and binding specific ligands, such as neurotransmitters or hormones, which activate the receptor and initiate signaling pathways within the cell. The extracellular N-terminus may also play a role in receptor activation and stabilization. The residues within 0.8 nm of the predicted ligand were extracted as the protein pocket. Utilizing AlphaFold-predicted GPCR structures offers a significant advantage by ensuring most data includes structural information, thereby increasing quantity of available data.

### Data preparation

To prepare the data, we first transformed the protein pocket into a graph representation, designating residues as nodes and contacting residue pairs as edges with a cutoff set as 0.5 nm. The protein pocket was defined based on a cutoff from the known ligand, retaining any residue whose atoms fell within this cutoff as part of the pocket Cutoff values were set at 0.6 nm for DeepGPCR_BC, in line with DeepBindGCN_BC^26^, and at 0.8 nm for DeepGPCR_RG, mirroring DeepBindGCN_RG^26^. Subsequently, we created a feature vector for each node using a 30-dimensional molecular vector trained using mol2vec ^27^. We used a parallel approach for the ligands, converting it into a graph representation with atoms as nodes and bonds as edges. We described each atom node with a one-hot-like representation, similar to the methodology implemented in GraphDTA ^16^.

### Training, test1 dataset, test2 dataset, and extra testing set

The data training, test1, test2, and extra test set are for DeepGPCR_BC and DeepGPCR_RG (**Table 1**).

**Table 1.** The number of samples in training and several test sets of DeepGPCR_BC/RG.

| DeepGPCR_BC | training datasets | test1 set | test2 set | Extra test set |
| --- | --- | --- | --- | --- |
|  | 35,000 | 3,531 | 1,668 | 9987 |
| DeepGPCR_RG | training datasets | test set |  | Extra test set |
|  | 186,000 | 8,408 |  | 51525 |

#### DeepGPCR_BC

Our study comprised 35,000 training datasets and 3,531 test1 sets, as shown in Figure S1. Although the protein-compound pairs in the test1 set were not included in the training dataset, the protein has been found in the training set, indicating that the test1 dataset is not fully independent. For a more accurate evaluation, 1,668 GPCR-compound pairs were chosen for the test2 set, ensuring full independence from the training dataset. All GPCRs lacking structural information were excluded from the training, test1, and test2 datasets. In the test1 set and validation, while the specific GPCR-ligand pairs were not found in the training, the GPCR may have formed pairs with other ligands. It is likely that the model has effectively learned this GPCR feature potentially leading to high accuracy predictions compared to those instances involving GPCR proteins not previously encountered in the training dataset. The test2 set posed a greater challenge, as not only GPCR-ligand pairs, but also the GPCRs themselves were not in the training set, mirroring real-world scenarios with novel GPCRs. However, predicting the structure and pocket of these novel GPCRs could significantly expand the size and diversity of the testing set. This would be particularly meaningful for applications, where the structure and pocket of GPCRs are often identified. We selected the 16 largest protein-related datasets to create an extra test set containing 9,987 protein-ligand pairs. We downloaded AlphaFold2 ^28^ predicted proteins from the AlphaFold Protein Structure Database (https://alphafold.ebi.ac.uk/). Ligand cofactors were modeled using COFACTOR ^29^. COFACTOR’s ligand-binding prediction involves identifying functional homologies from a non-redundant set of BioLiP templates^30^. Next, ligands are superposed to the predicted binding sites. Finally, the consensus binding sites are obtained by classified all ligands superposed to the query structure. In this work, we use COFACTOR within the local version of I-TASSER with the protein PDB file as the input. We then extracted residues within 0.6 nm of the selected ligand to form a pocket.

#### DeepGPCR_RG

For DeepGPCR_RG, we compiled 186,000 training datasets and 8,408 test sets (**Table 1)**. In analogue to DeepGPCR_BC, we selected the 16 largest protein-related datasets as an extra test set, comprising 51,525 protein-ligand pairs. The availability of Alphafold2 modeled structures, more abundant than GPCR structures from the PDB database, results in a larger number of GPCR proteins in the DeepGPCR_RG dataset compared to DeepGPCR_BC. Furthermore, unlike DeepGPCR_BC, DeepGPCR_RG does not exclude the protein-ligand pairs with binding affinities between 4 nM and 4000 nM. Consequently, the total data used to train DeepGPCR_RG is significantly larger, encompassing 186,000 pairs compared to DeepGPCR_BC’s which 35,000 (**Table 1)**.

### Model construction

Our model structure consists of two inputs,drug–target pair, and one output structure (**Figure 1A and C**). The ligand and pocket information are fed into two separate layers of a graph network. The outputs of the two graph networks are then merged into fully connected layers, culminating in a single node as the final output. For DeepGPCR_BC, the sigmoid activation function for binary prediction was employed, returning values between 0 and 1. Conversely, for DeepGPCR_RG, linear function for affinity prediction was implemented, yielding real values. The ReLU activation function was chosen as the activation function for each layer, except for the final node in the neural network. A dropout operation, with a rate of 0.2, was used following the pocket GCN layer, ligand GCN layer, and after the second merge layer.

**Figure 1.**
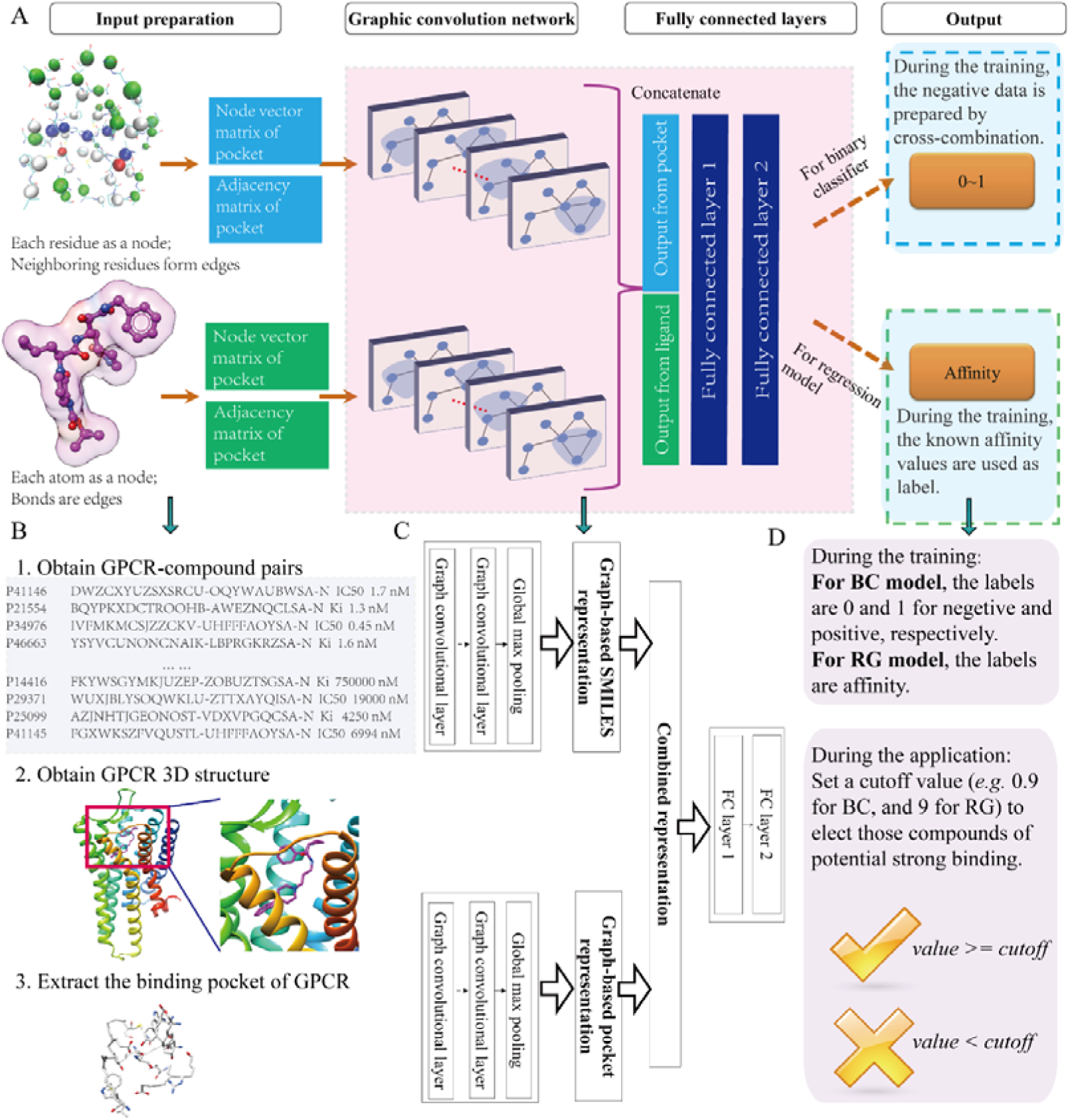
The workflow of DeepGPCR_BC and DeepGPCR_RG models construction and evaluation. A. illustrates the input graphic representation, model architecture, output, and performance evaluation metrics. B. Depicts the process of obtaining GPCR-compound pairs and a 3D binding pocket of GPCR. C. The detailed layer information of the GCN model. D. Briefly introduces the training label and the usage of the predicted label during application.

### Model training

The torch_geometric module generated input data and constructed the graph neural network. The input data was saved in PyTorch InMemoryDataset format. The PyTorch was operated for doing training. For DeepGPCR_BC, BCELoss (Binary Cross Entropy Loss) was selected as the loss function with the Adam optimizer. Similarly, for DeepGPCR_RG, MSEloss (Mean Squared Error Loss) was chosen as the loss function,also employing the Adam optimizer. The learning rate was 0.0005, and the total number of epochs was set to 2000. The model was saved at every 100-epoch interval. The final selection of epochs was based on the performance convergence observed in the test set. Performance metrics were recorded for the validation set after each epoch, facilitating the observation of model convergence and carrying different numbers of epoch-sensitive analyses.

### Virtual screening

We performed virtual screening against the ChemDiv database using the DeepGPCR_BC, DeepGPCR_RG and Schrödinger docking with Q9HC97 (GPR35) and GLP_1R serving as representative examples. The Chemdiv database, provided by ChemDiv company (https://www.chemdiv.com/), contains approximately1,500,000 compounds, most of which are purchasable from ChemDiv Company. For this purpose, GPR35 sequence were sourced from UniProt, and 3D structure models were constructed using AlphaFold2. The binding sites were predicted using COFACTOR, with the selection of the binding site candidate nearest to the N-terminal of the protein. The structure of GLP_1R was from PDB structure (PDBID:7s15^31^), the binding site was determined by the known ligand 82L within the PDB structure. We used the trained models as the core component during the screening and developed custom scripts. For each protein, multiple binding site candidates were considered, and the screening process utilizing the selected binding site. The ChemDiv database was screened using the DeepGPCR_BC, DeepGPCR_RG, and Schrödinger docking. The Schrödinger docking procedures are outlined in Supplementary materials section 1.

### Performance metrics for binary classification model

To assess model performance, we employed various evaluation metrics including AUC (Area Under the ROC Curve), TPR (True Positive Rate), Precision, Accuracy, MCC (Matthews Correlation Coefficient), and F1 score. AUC, representing the area under the ROC (Receiver Operating Characteristic) curve, ranges from 0.5 to 1, where 0.5 denotes a random classifier, and 1 indicates a perfect classifier. TPR, known as recall, refers to the proportion of true positive instances that are correctly predicted as positive. Precision measures the ratio of true positives to the sum of true positives and false positives, representing the proportion of true positives to the total predicted positives by the model. Accuracy, ranging from 0 to 1, is the ratio of correctly classified samples out of the total number of samples. MCC, a correlation coefficient, quantifies the relationship between the actual and predicted binary classifications. It takes values between -1 to 1. A value of -1 indicates a perfect negative correlation, 1 reveals a perfect positive correlation, and 0 indicates no correlation. The F1 score, an essential metric for binary classification models, combines Precision, and Recall to provide a balanced perspective on the performance of model.

It should be noted that the aforementioned performance metrics are tailored to evaluate binary classification prediction ranging from 0 to1.In contrast, Schrödinger and Autodock Vina predicted value of linear; hence, we used -6 kcal/mol as the cutoff, those scores greater than -6 kcal/mol was assigned value 0, indicating non-bind, and those scores equal to or less than -6 kcal/mol were assigned a value of 1, representing binding capability. This approach enables the evaluation of performance with the above evaluation metrics.

### Performance metrics for the Regression model

We used RMSE (Root Mean Squared Error), MSE (Mean Squared Error), Pearson (Pearson correlation coefficient), Spearman (Spearman correlation coefficient) and C-index (Concordance Index, CI) to evaluate the performance of the regression model. RMSE is computed as the square root of the average squared differences between the predicted and actual values. MSE is the average of the squared differences. The Pearson correlation measures the linear relationship between these predicted and actual values. Spearman correlation is a non-parametric assessment of the monotonic relationship between variables. C-index measures the ability of a model to rank the observed outcomes correctly in terms of their relative risk or event occurrence probabilities.

### Compounds and reagents

All compounds for GPR35 and GLP-1R were obtained from TOPSCIENCE (Shanghai, China). Zaprisnast was purchased from Sigma Aldrich (Shanghai, China). Taspoglutide was purchased from MedChem Express (Shanghai, China). Hank’s balanced salt solution (HBSS), HEPES, fetal bovine Serum (FBS), penicillin, streptomycin and F12 medium were obtained from Invitrogen (Shanghai, China).

### Cell culture

CHO-GPR35 cells were the same as our previously reported^32^. CHO-GPR35 cells were cultured in an F12 medium supplemented with 10% FBS, penicillin (50 μg/mL), streptomycin (100 μg/mL), and zeocin (200 μg/mL) at 37◦C under 5% CO_2_. HEK293 cells were purchased from the National Collection of Authenticated Cell Cultures (Shanghai, China) and cultured in DMEM medium supplemented with 10% FBS, penicillin (50 μg/mL) and streptomycin (100 μg/mL) and at 37◦C under 5% CO_2_.

### Construction of HEK293-GLP-1R stable cells

HEK293 cells were transfected with 8 μg of pcDNA3.1-GLP-1R plasmid mixed with 24 μL of Lipofectamine 2000 reagent (Invitrogen). After 24 hours post-transfection, clones were selected using a complete medium containing 600 μg/mL G418 (LabLEAD Co., Beijing, China) and 6 μg/mL blasticidin S (Beyotime Co., Shanghai, China). Stable clones were selected through treatment with 600 μg/mL G418 and 6 μg/mL blasticidin S for 3-4 weeks to obtain the successfully transfected cell line HEK293-GLP-1R. Following culture for 3-4 months, the stably transfected cell line HEK293-GLP-1R was established.

### Dynamic mass redistribution (DMR) assay

When the cells approached 90% confluence, they were seeded in 384 well biosensor plates with a density of 1.5×10^4^ CHO-GPR35 or 2.5×10^4^ HEK293-GLP-1R cells/well and cultured for 24 h. The culture medium in the 384-well biosensor plates was replaced with 30 μL of Hank’s balanced salt solution (1×HBSS) and then further incubated inside the system for 1 h before measurement. For the DMR agonism assay, a 2-min baseline was first established, followed by adding compounds using the multi-channel pipette, and the compound-triggered DMR responses were recorded for approximately 1 h. Subsequently, the baseline was re-established, Zaprinast (a known GPR35 agonist) at a fixed concentration (100 nM) was added and the DMR responses induced were recorded for 1 h, while taspoglutide (a known GLP-1R agonist) at a fixed concentration (1 μM) was added, and the DMR responses induced were recorded for 90 min. For DMR antagonism assay, cells were initially treated with either an antagonist, or compound for 1 h in GPR35 and 90 min in GLP-1R assays, respectively. Afterwards, the baseline was re-established, followed by adding Zaprinast at a fixed concentration (100 nM) and taspoglutide at a fixed concentration (1 μM), and then monitoring the DMR responses induced by zaprinast for 1 h and taspoglutide for 90 min. All DMR responses were background corrected.

## Results and discussion

The construction workflow of our two models is illustrated in **Figure 1**. We represented pockets and ligands as graph representations for input and employed the Graph Convolutional Network along with fully connected layers to train the model (**Figure 1B**). The binary classifier model generates output values ranging from 0 to 1, wherein values closer to 0 signifying weak or no binding, while values closer to 1 denoting strong binding interactions. For the regression model, the output is a numerical value, where larger values correspond to stronger binding interactions. For performance evaluation, we employed 16 GPCR related datasets as an extra test set. However, due to the unavailability of experimental 3D structures and binding pocket information for these GPCRs, we resorted to obtaining the AlphaFold predicted 3D structures ^33^ and utilizing COFACTOR ^34^ to predict the ligand binding pocket (**Figure 1B**).

### Performance of DeepGPCR_BC model

We assessed the effectiveness of our DeepGPCR_BC model on both the training and testing datasets and summarized the results in **Table S1** and **Figure S2**. Our findings revealed that the model’s performance converged and remained relatively stable after approximately 100 epochs. Our model demonstrated excellent performance on the test set1 with an AUC of 0.97, TPR of 0.90, Precision of 0.90, Accuracy of 0.92, and MCC of 0.84 at epoch 2000.

However, the high performance on the test1 dataset may be due to the inclusion of its protein in the training set, implying that the model had fully learned its pocket feature during training. Therefore, this dataset is not entirely independent of the training set. To assess the model’s ability to generalize the non-trained protein pockets, its performance on the test2 dataset was evaluated. The performance metrics in test set2 at epoch 2000 revealed a decrease, with AUC, TPR, Precision, Accuracy, and MCC, dropping to 0.72, 0.46, 0.46, 0.70, and 0.16 respectively. (**Table S2**). Since the test set2 only contains one protein (P29274), it provides a limited scope for fully evaluating the model’s reliability, we afterwards test the performance on a more diversified and fully independent extra dataset.

The selected proteins with the predicted pockets, and the performance are shown in **Figure 2** and **Table 2, representative**. The model achieved metrics 0.72, 0.73, 0.73, 0.68, 0.37, and 0.64 for all the datasets for AUC, TPR, Precision, accuracy, MCC, and F1 score, respectively. Upon evaluating individual protein performance, the model demonstrated optimal results for P20309 and P35372. Notably, DeepGPCR_BC outperforms the Schrödinger’s Glide and AutoDock Vina docking as detailed in **Table 3** and **S3**. It should be noted that the DeepGPCR_BC score is the output score with a range between 0 to 1. In contrast, the Schrödinger and Autodock Vina score represent the predicted binding affinity of the ligand to the receptor in a continuous unit, kcal/mol. Here we set -6 kcal/mol as the cutoff distance, the values larger than -6 kcal/mol were assigned a value of 0, indicating non-bind, and those scores less than and equal to -6 kcal/mol were given a value of 1, indicating able to bind. In this way, we can evaluate their performance with evaluation metrics of AUC, TPR, precision, accuracy, and MCC. The Schrödinger dock yielded poor performance, with an AUC of 0.45, TPR of 0.29, Precision of 0.43, accuracy of 0.45, MCC of -0.10, and F1 score of 0.35 across all datasets. The poor performance of the Schrödinger dock could be attributed to the use of generalized protein datasets primarily comprising soluble proteins, which have significantly different physical-chemical properties than GPCR proteins. Similarly, Autodock Vina also exhibited poor performance, with an AUC of 0.49, TPR of 0.78, Precision of 0.50, accuracy of 0.49, MCC of -0.02, and F1 score of 0.61 for the entire dataset (**Table S3**).

**Figure 2.**
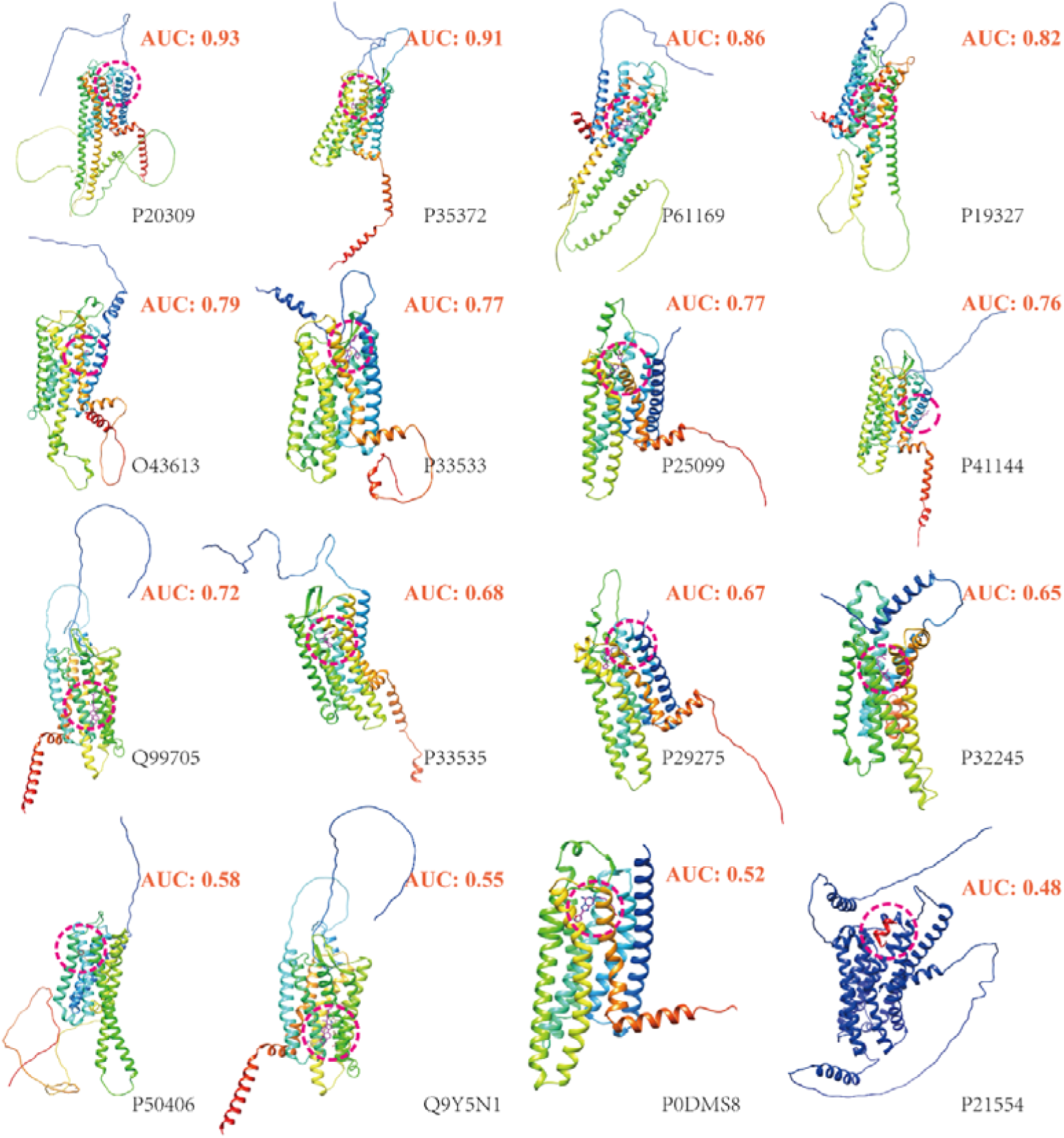
The modeled protein for the extra test set and the modeled ligands determines the pockets. Out of the 16 test cases, 6 achieved an AUC larger than 0.7, and 10 achieved an AUC larger than 0.6. It should be noted that none of the 16 proteins were included in the training set.

**Table 2.** DeepGPCR_BC performance on an extra dataset with modeled GPCR protein and predicted pocket.

| Name | AUC | TPR | precision | accuracy | MCC | F1<br>score | pos_num | neg_num |
| --- | --- | --- | --- | --- | --- | --- | --- | --- |
| P20309 | 0.93 | 0.88 | 0.92 | 0.88 | 0.76 | 0.90 | 664 | 478 |
| P35372 | 0.91 | 0.84 | 0.86 | 0.85 | 0.71 | 0.85 | 1054 | 1077 |
| P61169 | 0.86 | 0.85 | 0.69 | 0.79 | 0.58 | 0.76 | 529 | 805 |
| P19327 | 0.82 | 0.77 | 0.86 | 0.76 | 0.49 | 0.81 | 948 | 451 |
| O43613 | 0.79 | 0.65 | 0.49 | 0.76 | 0.41 | 0.56 | 226 | 734 |
| P33533 | 0.77 | 0.84 | 0.75 | 0.73 | 0.42 | 0.79 | 609 | 399 |
| P25099 | 0.77 | 0.47 | 0.51 | 0.81 | 0.37 | 0.49 | 199 | 813 |
| P41144 | 0.76 | 0.84 | 0.74 | 0.72 | 0.39 | 0.79 | 552 | 352 |
| Q99705 | 0.72 | 0.48 | 0.71 | 0.65 | 0.31 | 0.57 | 450 | 483 |
| P33535 | 0.68 | 0.72 | 0.75 | 0.65 | 0.23 | 0.74 | 913 | 443 |
| P29275 | 0.67 | 0.19 | 0.23 | 0.77 | 0.07 | 0.21 | 147 | 794 |
| P32245 | 0.65 | 0.40 | 0.57 | 0.63 | 0.20 | 0.47 | 404 | 575 |
| P50406 | 0.58 | 0.22 | 0.87 | 0.41 | 0.15 | 0.35 | 913 | 339 |
| Q9Y5N1 | 0.55 | 0.25 | 0.89 | 0.32 | 0.03 | 0.39 | 1308 | 201 |
| P0DMS8 | 0.52 | 0.16 | 0.55 | 0.60 | 0.10 | 0.25 | 566 | 805 |
| P21554 | 0.48 | 0.30 | 0.36 | 0.61 | 0.05 | 0.32 | 571 | 1238 |
| ALL | 0.72 | 0.73 | 0.73 | 0.68 | 0.37 | 0.64 | 10053 | 9987 |

**Table 3.** The Schrödinger docking performance on an extra dataset with modeled GPCR protein and predicted pocket. We used -6 Kcal/mol as the cutoff, those scores > -6 Kcal/mol was assigned a value of 0 (indicating non-bind), and those scores ≤ -6 Kcal/mol were assigned a value of 1 (indicating able to bind).

| Name | AUC | TPR | precision | accuracy | MCC | F1<br>score | pos_num | neg_num |
| --- | --- | --- | --- | --- | --- | --- | --- | --- |
| P29275 | 0.56 | 0.85 | 0.18 | 0.36 | 0.10 | 0.29 | 147 | 794 |
| P19327 | 0.56 | 0.20 | 0.84 | 0.43 | 0.15 | 0.32 | 948 | 451 |
| P20309 | 0.52 | 0.25 | 0.63 | 0.48 | 0.05 | 0.36 | 664 | 478 |
| Q99705 | 0.51 | 0.02 | 1.00 | 0.53 | 0.10 | 0.03 | 450 | 483 |
| P25099 | 0.50 | 0.97 | 0.20 | 0.22 | 0.01 | 0.33 | 199 | 813 |
| P41144 | 0.50 | 0.00 | 1.00 | 0.39 | 0.03 | 0.00 | 552 | 352 |
| Q9Y5N1 | 0.50 | 0.00 | 1.00 | 0.13 | 0.01 | 0.00 | 1308 | 201 |
| O43613 | 0.50 | 0.02 | 0.25 | 0.76 | 0.00 | 0.03 | 226 | 734 |
| P33533 | 0.50 | 0.00 | 0.00 | 0.39 | -0.07 | 0.00 | 609 | 399 |
| P61169 | 0.49 | 0.00 | 0.08 | 0.60 | -0.06 | 0.00 | 529 | 805 |
| P0DMS8 | 0.49 | 0.92 | 0.41 | 0.42 | -0.02 | 0.57 | 566 | 805 |
| P33535 | 0.49 | 0.78 | 0.67 | 0.59 | -0.02 | 0.72 | 913 | 443 |
| P21554 | 0.48 | 0.02 | 0.16 | 0.65 | -0.07 | 0.04 | 571 | 1238 |
| P35372 | 0.45 | 0.74 | 0.46 | 0.45 | -0.12 | 0.57 | 1054 | 1077 |
| P50406 | 0.44 | 0.20 | 0.63 | 0.33 | -0.12 | 0.31 | 913 | 339 |
| P32245 | 0.42 | 0.00 | 0.01 | 0.49 | -0.27 | 0.00 | 404 | 575 |
| ALL | 0.45 | 0.29 | 0.43 | 0.45 | -0.10 | 0.35 | 10053 | 9987 |

### Performance of DeepGPCR_RG model

The performance of the DeepGPCR_RG model across different training epochs is detailed in **Table S4**. The training was performed over 2000 iterations, with performance metrics including RMSE, MSE, Pearson’s correlation coefficient and Spearman’s rank correlation coefficient recorded at intervals of every 200 epochs.

Regarding the training data, a significant improvement in performance was observed between 200^th^ and 2000^th^ epoch. The RMSE decreased from 0.72 to 0.64, showing a reduction in the variance of the prediction errors. The Pearson correlation showed a slight increase from 0.84 to 0.87, signifying a strong linear relationship between the actual and predicted outputs. The performance on the test data remained relatively stable across different epochs (Table S4). Indicating that although the model’s performance on training data improved with successive epochs, its ability to generalize to unseen data did not significantly improve over the observed period. Furthermore, DeepGPCR_RG’s performance was assessed on an additional test set composed of data. This model’s performance on the additional test set was subsequently compared to that of Schrödinger and Vina docking methods on the same datasets. Performance measures for DeepGPCR_RG on the extra test datasets is depicted in **Table 4**. Overall, DeepGPCR_RG showed average RMS error (RMSE) and Mean Square Error (MSE) values of 1.34 and 1.80 respectively. Pearson’s and Spearman’s correlation coefficients averaged at 0.39 and 0.35, respectively, across all datasets. This suggests a moderate monotonic and a linear relationship between actual and predicted outputs.

**Table 4.** The DeepGPCR_RG performance on regression model’s extra test datasets.

| **Name** | **rmse** | **mse** | **pearson** | **spearman** | **total_num** |
| --- | --- | --- | --- | --- | --- |
| P20309 | 1.35 | 1.82 | 0.65 | 0.69 | 2006 |
| P35372 | 1.19 | 1.41 | 0.57 | 0.56 | 4236 |
| P25099 | 1.19 | 1.42 | 0.49 | 0.48 | 2932 |
| P33533 | 1.35 | 1.82 | 0.48 | 0.47 | 1969 |
| P41144 | 1.31 | 1.71 | 0.46 | 0.44 | 2173 |
| P32245 | 1.37 | 1.88 | 0.42 | 0.37 | 2613 |
| P61169 | 1.16 | 1.35 | 0.42 | 0.41 | 4460 |
| P33535 | 1.53 | 2.36 | 0.38 | 0.37 | 2962 |
| O43613 | 1.22 | 1.49 | 0.35 | 0.33 | 3811 |
| Q9Y5N1 | 1.34 | 1.80 | 0.34 | 0.32 | 3693 |
| Q99705 | 1.25 | 1.56 | 0.31 | 0.25 | 3610 |
| P19327 | 1.26 | 1.59 | 0.29 | 0.29 | 4632 |
| P0DMS8 | 1.46 | 2.13 | 0.26 | 0.27 | 3797 |
| P29275 | 1.26 | 1.59 | 0.18 | 0.14 | 1804 |
| P50406 | 1.62 | 2.63 | 0.12 | 0.09 | 3297 |
| P21554 | 1.59 | 2.53 | 0.11 | 0.10 | 3530 |
| Average | 1.34 | 1.80 | 0.39 | 0.35 | 51525 |

In contrast, Schrödinger’s docking method showed higher average RMSE and MSE values, illustrating a larger discrepancy between the predicted and actual values. Average Pearson and Spearman correlation coefficients were both negative -0.04, in **Table S5**, suggesting that the Schrödinger’s ability in predicting dataset values was weaker than to DeepGPCR_RG.

Lastly, the performance of the Vina docking method as shown in **Table S6**, also demonstrated higher average RMSE and MSE values. Like Schrödinger’s docking method, both Pearson’s and Spearman’s correlation coefficients are negative (average -0.06 and -0.07 respectively).

In summary, on the additional test datasets, DeepGPCR_RG was superior to the other two methods, demonstrating lower error rates and better correlation coefficients. This illustrates DeepGPCR_RG enhanced prediction accuracy and its stronger alignment with actual values compared to Schrödinger and Vina docking methods

### Screening against target Q9HC97 (GPR35) by DeepGPCR_BC, DeepGCPR_RG, and Schrödinger

Q9HC97 was chosen to demonstrate the applications of our models in screening potential therapeutic compounds. GPR35 has been identified as a potential target for various diseases ^35^. The screening procedures are shown in **Figure 3**. Using our DeepGPCR_BC and DeepGPCR_RG models, we screened 102,592 candidates with a DeepGPCR_BC score≥ 0.999 and a DeepGPCR_RG score ≥9 for Q9HC97. We also calculated the Schrödinger score for those candidates. The DeepGPCR_BC score ≥ 0.999, DeepGPCR_RG≥9, and Schrödinger score≤-6.7 Kcal/mol (**Table 5**), and those candidates are selected for final experimental validation.

**Figure 3.**
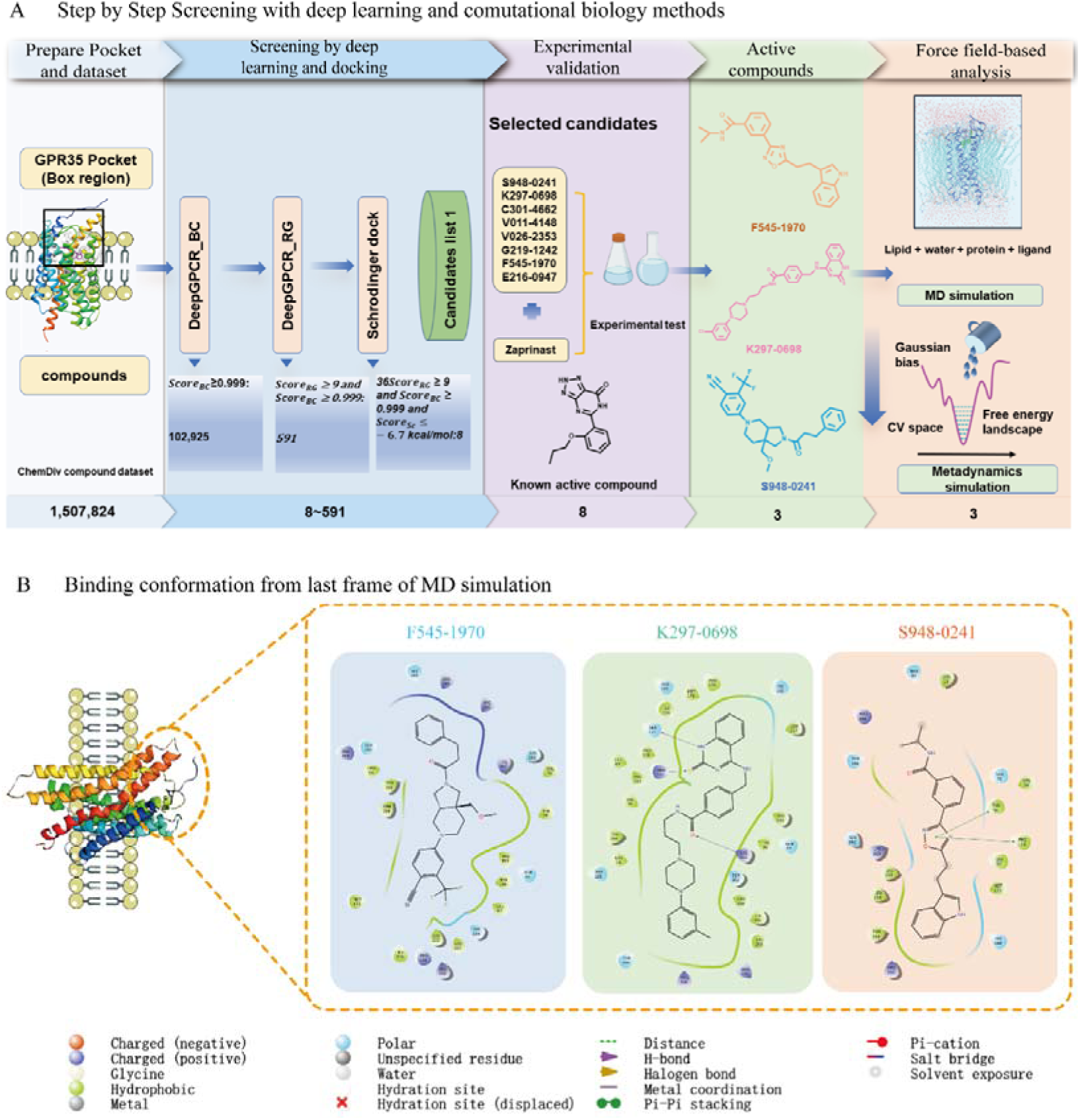
GPR35 screening pipeline and Identification of Active Compounds. A. Schematic representation of the stepwise screening process leading to the discovery of 3 active molecules. B. conformation of last frame from the MD simulation and 2D protein-ligand interaction diagrams for the identified active compounds.

**Table 5.** The candidate list of GPR35 from screening with DeepGPCR_BC score ≥ 0.999, DeepGPCR_RG≥9, and Schrödinger score≤ -6.7 Kcal/mol.

| Chemdiv ID | DeepGPCR_BC | DeepGPCR_RG | Schrödinger<br>(Kcal/mol) |
| --- | --- | --- | --- |
| S948-0241 | 1.00 | 9.39 | -7.61 |
| K297-0698 | 1.00 | 9.08 | -7.37 |
| C301-4662 | 1.00 | 9.11 | -7.19 |
| V011-4148 | 1.00 | 9.76 | -7.06 |
| V026-2353 | 1.00 | 9.26 | -6.78 |
| G219-1242 | 1.00 | 9.56 | -6.74 |
| F545-1970 | 1.00 | 9.01 | -6.74 |
| E216-0947 | 1.00 | 9.25 | -6.74 |

To further investigate the reliability of using DeepGPCR_RG and Schrödinger’s software independently, we conducted experimental validation over another candidate list by using DeepGPCR_RG and Schrödinger (DeepGPCR_RG≥10, Schrödinger score≤-6.35 Kcal/mol) (**Table S7**). Together with Table 5, a total of 12 candidates were forwarded for experimental validation, the structures of those compounds are shown in **Figure S3**.

### Screening against target GLP-1R by DeepGPCR_BC, DeepGCPR_RG, and Schrödinger

We selected GLP-1R to further validate our model’s capability in screening potential therapeutic compounds. GLP-1R, identified as a potential target for various diseases including cancer, was our focus. The screening procedures are shown in **Figure S4**. Using our DeepGPCR_BC and DeepGPCR_RG models, we screened 158 candidates with a DeepGPCR_BC score≥ 0.999 and a DeepGPCR_RG score ≥9.5 for **GLP-1R**. We also calculated Schrödinger score for these candidates. Selection criteria were set at a DeepGPCR_BC score ≥ 0.999, DeepGPCR_RG≥9.5, and Schrödinger score≤-8.7 Kcal/mol (**Table 6**). For further investigation, the standalone reliability of DeepGPCR_RG and Schrödinger’s software,, we select 3 based on a DeepGPCR_RG ≥10, Schrödinger score≤-6.35 kcal/mol) (**Table S8**). Combining the results from with

**Table 6.** The candidate list of GLP_1R from screening with DeepGPCR_BC score≥ 0.999, DeepGPCR_RG≥9.5, and Schrödinger score≤ -8.7 Kcal/mol.

| **Chemdiv ID** | **DeepGPCR_RG** | **DeepGPCR_BC** | **Schrödinger** |
| --- | --- | --- | --- |
| V027-3795 | 9.68 | 1.00 | -9.14 |
| V012-1447 | 9.53 | 1.00 | -9.52 |
| V009-0856 | 9.55 | 1.00 | -8.82 |
| V005-2405 | 9.55 | 1.00 | -8.80 |
| S978-0648 | 9.70 | 1.00 | -8.82 |
| C875-0922 | 9.56 | 1.00 | -9.00 |
| G818-0282 | 9.73 | 1.00 | -9.03 |
| S954-5266 | 9.67 | 1.00 | -9.11 |
| C700-1841 | 9.56 | 1.00 | -9.45 |
| SB04-0468 | 9.56 | 1.00 | -9.58 |
| Y043-3164 | 9.76 | 1.00 | -8.83 |
| T158-1384 | 9.64 | 1.00 | -8.89 |

Table 6, a total of 15 candidates were selected for experimental validation, the structures of those compounds shown in **Figure S5**.

### Characteristics of selective candidates on GPR35

We applied the GPR35 overexpressed CHO cells to assess the activity of selective candidates as well as Zaprinast, an endogenous ligand of GPR35. Among the 12 candidates, S948-0241, K297-0698 and F545-1970 exhibit desensitization effects on the GPR35 receptor when stimulated by the agonist Zaprinast, and they do not show a significant increase in DMR signal in CHO-GPR35 cells, indicating that they possess GPR35 receptor antagonistic activity, with relatively weak activity and IC50 values around 30-80 μM (**Figure 4 A-D**). E014-0043, C301-4662, V026-2353, and G219-1242 only exhibit desensitization effects on the GPR35 receptor when stimulated at high concentrations with poor desensitization effects, and they do not show a significant increase in DMR signal in CHO-GPR35 cells, indicating that they have weak GPR35 receptor antagonistic activity (**Figure 4 E-H**). E146-0380, D103-0816, L311-0042, V011-4148, and E216-0947 do not exhibit desensitization effects or only show weak desensitization effects when stimulated by the agonist Zaprinast on the GPR35 receptor, indicating an absence of GPR35 receptor activity (**Figure 4 I-K**). These results suggest that 3 out of 12 candidates possess significant selective antagonistic activity targeting GPR35.

**Figure 4.**
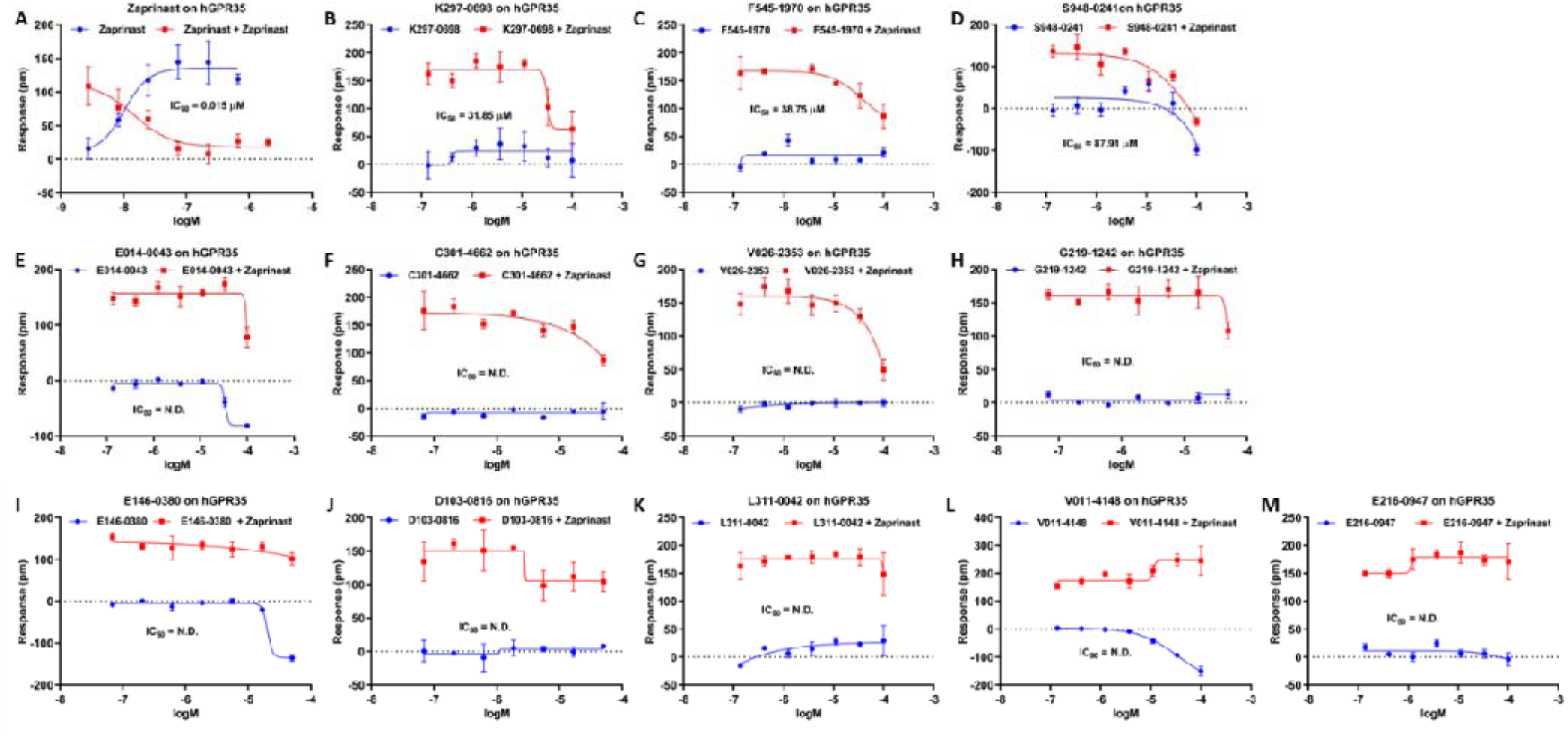
Characteristics of selective candidates on GPR35. (A) Zaprinast, an endogenous ligand of GPR35 and (B-K) selective candidates of GPR35. N.D. denotes not determined.

### Activity validation of selective candidates on GLP-1R

Next, we utilized the GLP-1R overexpressed HEK293 to test the activity of selective candidates for GLP-1R, along with Taspoglutide, a positive control as a GLP-1 agonist. Among the 15 selective candidates, V005-2405 exhibits desensitization effects on the GLP-1R receptor when stimulated by the agonist Taspoglutide. This effect is dose-dependent, with an IC50 value of 9.60 μM (**Figure 5 A and B**). Additionally, V005-2405 does not induce a DMR signal, indicating that V005-2405 possesses GLP1R receptor antagonistic activity. C700-1841 and G764-0921 exhibit desensitization effects on GLP-1R when stimulated by the agonist Taspoglutide (partial desensitization, approximately 40% inhibition) (**Figure 5 C and D**). This effect is dose-dependent, and C700-1841 and G764-0921 do not induce a DMR signal, indicating that they possess partial GLP-1R antagonistic activity. S954-5266 only exhibit desensitization effects on GLP-1R when exposed to high concentrations at 200 μM of the agonist Taspoglutide. At other concentrations, it does not inhibit Taspoglutide activity and induce a DMR signal. This suggests that S954-5266 has GLP1R receptor antagonistic activity at 200 μM (**Figure 5 E**). V009-0856 and V027-3795 exhibit desensitization effects on GLP-1R when exposed to high concentration with 200 μM of the agonist Taspoglutide. However, they either do not induce a DMR signal or exhibit a weak signal in GLP-1R-HEK293 cells, indicating that these two compounds have GLP-1R receptor antagonistic activity at 200 μM (**Figure 5 F and G**). The remaining compounds either do not have GLP-1R activity or exhibit weak GLP-1R activity (**Figure 5 H-P**). These findings indicate that 6 out of the 15 candidates exhibit noteworthy selective antagonistic activity against GLP-1R.

**Figure 5.**
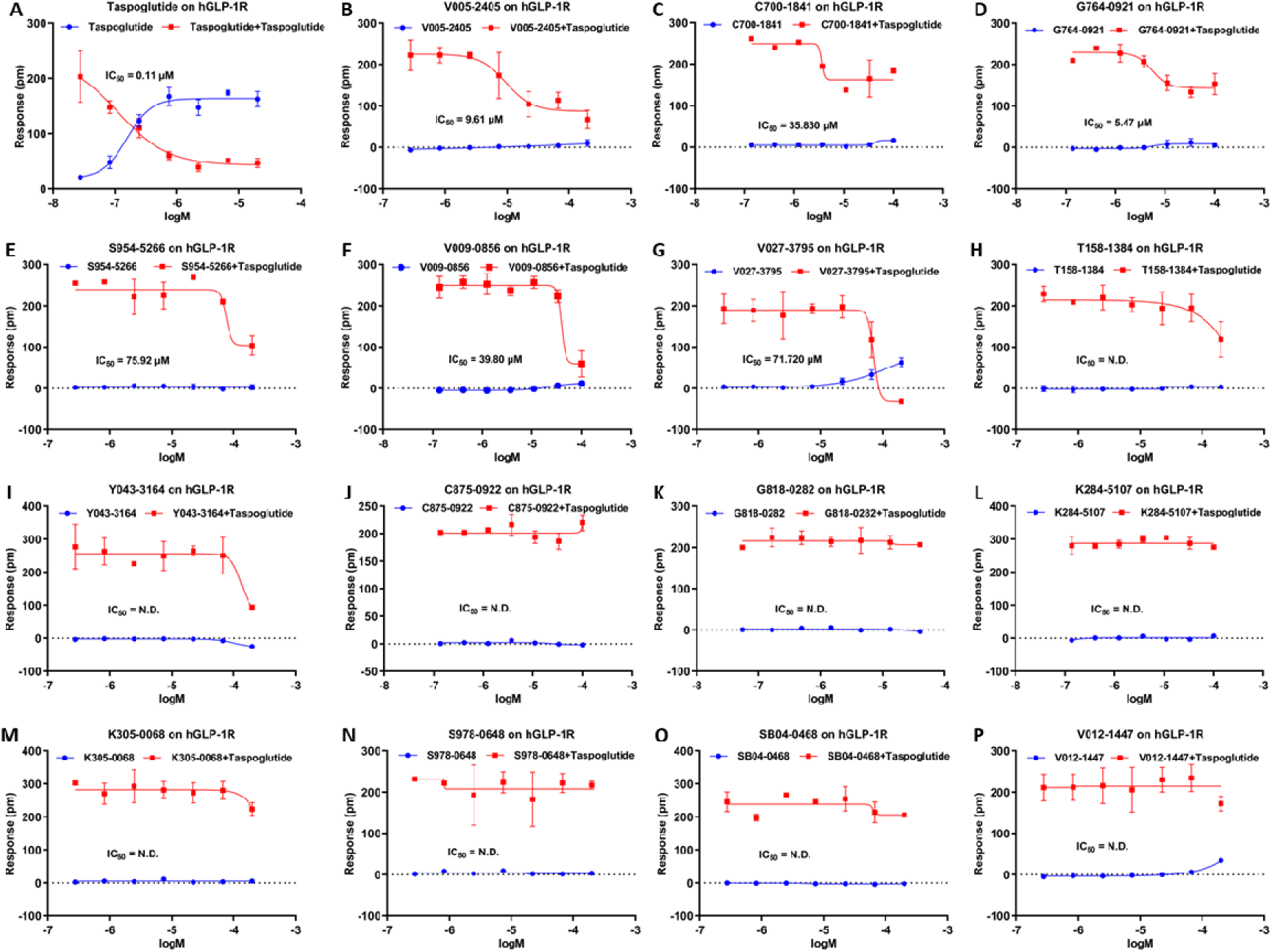
Characteristics of selective candidates on GLP-1R. (A) Taspoglutideis a former experimental drug, a glucagon-like peptide-1 agonist (GLP-1 agonist) and (B-K) selective candidates of GLP-1R. N.D. denotes not determined.

### Detailed analysis of the GPR35 with those identified active compounds

In **Figure 6**, we present the docking interactions of GPR35 with three active molecules and a known active control molecule in both 3D and 2D representations. The interaction between GPR35 and K297-0698, as shown in **Figure 6A**. The primary interactions between the control compounds and proteins are characterized by hydrophobic, electrostatic, and polar forces. TYR259, LEU97, and PRO176 predominantly engage in hydrophobic interactions with the cyclohexane moiety of compound K297-0698. The positive charged ARG151 establishes a robust π-cation interaction with the pyrimidine ring of compound K297-0698, while the charged interaction between ARG240 and the chlorine atom of the compound facilitates the formation of hydrogen bonds between the oxygen atom in the structure of PHE163 and the oxygen atom of the amide bond in compound K297-0698.

**Figure 6.**
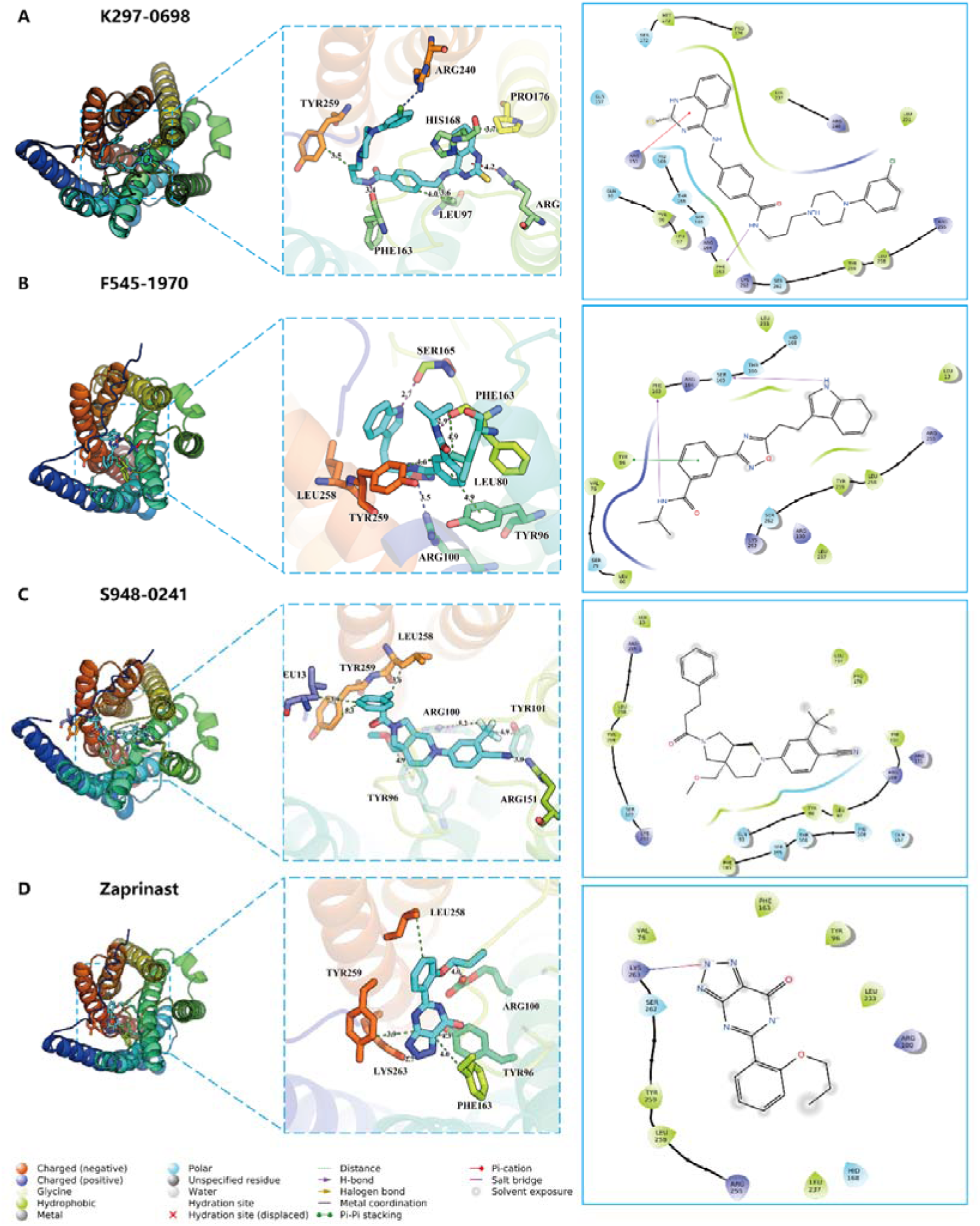
the docking interactions of GPR35 with three active molecules and a known active control molecule in both 3D and 2D representations. A. Residue-specific interactions between pocket residues and compound K297-0698. Interaction diagram of pocket residues with compound K297-0698. B. Residue-specific interactions between pocket residues and compound F545-1970. Interaction diagram of pocket residues with compound F545-1970. C. Residue-specific interactions between pocket residues and compound S948-0241. Interaction diagram of pocket residues with compound S948-0241. D. Residue-specific interactions between pocket residues and known active compound Zaprinast. For all the plots, Residues that mainly provide hydrophobic, charged, π-cation and polar interactions are colored in green, blue,red and purple.. The coloration of proteins is determined by the B-factor (thermal motion) of carbon atoms, exhibiting a gradient that transitions from orange to yellow, then green, blue, and ultimately culminating in purple. Residues that mainly provide hydrophobic, electrostatic and polar interactions are colored in green, blue and purple.

The interactions between GPR35 and F545-1970 are shown in **Figure 6B**. The control compounds and proteins predominantly engage in hydrophobic, electrostatic, and polar interactions. LEU258, TYR259, LEU80, and TYR96 are primarily involved in hydrophobic interactions with the cyclohexane portion of compound F545-1970. Notably, TYR96 establishes π-π interactions with the cyclohexane ring of the compound. ARG100 and the 1,2,4-oxydiazole moiety of compound F545-1970 engage in an electrostatic interaction. Additionally, the oxygen atom in the structure of SER165 and the phenyl group of PHE163 form hydrogen bonds with the nitrogen atom of compound F545-1970.

The interactions between GPR35 and S948-0241 are shown in **Figure 6C**. The primary interactions between the control compounds and proteins are hydrodynamic and electrostatic in nature. LEU258, TYR259, LEU13, and TYR96 predominantly engage in hydrophobic interactions with the cyclohexane moiety of compound S948-0241, with TYR96 specifically forming π-alkyl interactions with the methylene groups of the compound. Furthermore, ARG100 and ARG151 establish electrostatic interactions with the positively charged elements of the trifluoromethyl group in the compound, namely the fluorine atoms and the nitrogen atom of the cyanide group.

The interaction between GPR35 and known active compounds Zaprinast are shown in **Figure 6D**. The control compounds and proteins predominantly engage in hydrodynamic and electrostatic interactions. LEU258, TYR259, PHE163, and TRP96 primarily establish hydrophobic interactions with the carbon atoms in the backbone of the control compounds. Conversely, LYS263 and ARG100 are involved in electrostatic interactions with the framework of the control compounds. Additionally, LYS236 engages in a salt bridge interaction with the nitrogen atom located at position 2 of the triazole.

### Detailed analysis of the GLP-R1 with those identified active compounds

In **Figure 7**, we present the docking interactions of GLP-1R pocket with three active molecules in detailed 3D representations. The overall binding view and 2D interaction plot can be found in **Figure S6**.

**Figure 7.**
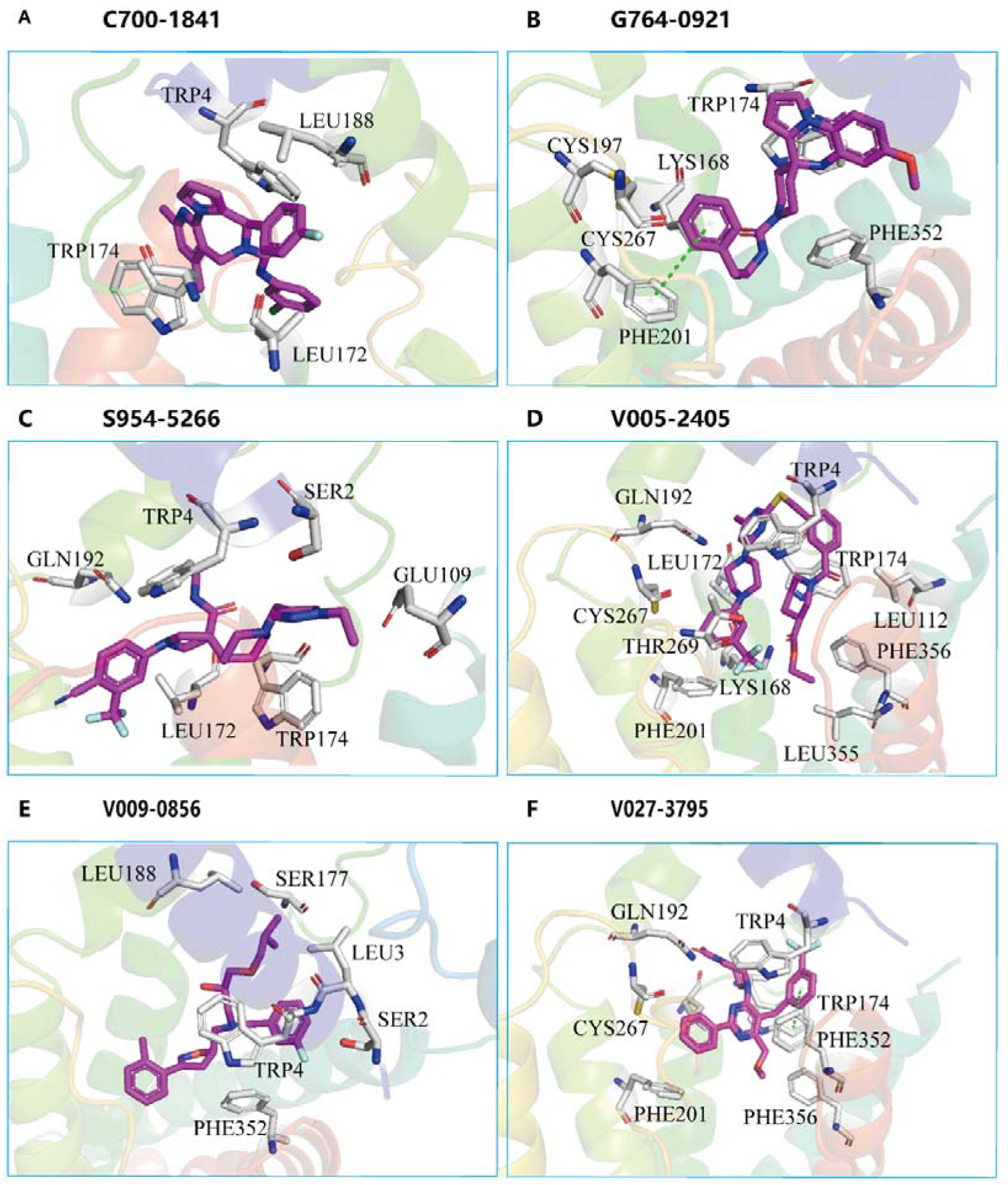
the predicted interactions of GLP_R1 with six active molecules from docking. A, Residue-specific interactions between pocket residues and compound C700-1841. B. Residue-specific interactions between pocket residues and compound G764-0921. C. Residue-specific interactions between pocket residues and compound S954-5266. D. Residue-specific interactions between pocket residues and known active compound V005-2405. E. Residue-specific interactions between pocket residues and known active compound V009-0856. F. Residue-specific interactions between pocket residues and known active compound V027-3795.For all the plots, Residues that mainly provide hydrophobic, charged, π-cation and polar interactions are colored in green, blue,red and purple. The coloration of proteins uses pymol’s rainbow.

The interactions between GLP-1R and C700-1841 are shown in **Figure 7A**, which shows hydrophobic are dominant. Hydrophobic residues, including TRP4, LEU188, TRP174, LEU172 are closely contacted with the C700-1841. The interaction between GLP-1R and G764-0921 are shown in **Figure 7B**. The compound and protein predominantly engage in hydrophobic, electrostatic, and polar interactions. The TRP174 PHE352 and PHE201are primarily involved in hydrophobic interactions. Notably, PHE201 establishes π-π interactions with the benzene like ring of the compound. Polar and charge residues CYS197 and LYS168 also have electrostatic and polar interaction with the compounds. The interactions between GLP_1R and S954-5266 are shown in **Figure 7C**, residues GLN192, LEU172, TRP 174, TRP 4, SER2, and GLU109 have close contact. Consequently, the interaction between the compound and the protein is structurally characterized by hydration.

The V005-2405 contacted with GLP-1R significantly, due to its larger size, shown in **Figure 7D**, have larger interactions with many hydrophobic residues such as LEU TRP and PHE. Some polar residue GLN192, CYS267, THR269 are also contributed to the binding interactions. The charge residue LYS168 also may have interaction with the carbon atom of trifluoromethyl of the compound.

The V009-0856 have formed hydrophobic and polar interaction with GLP_1R, shown in **Figure 7E**. The hydrophobic Residues, such as LEU3, TRP4, and PHE352. The polar residue SER2 is in close contact with the compound and may form Hydrogen Bonding with the fluorine atom on the benzene ring of the compound.

The V027-3795 have formed hydrophobic and polar interaction with GLP-1R, shown in **Figure 7F**. The Residues containing a benzene ring, such as TRP4, TRP174, PHE201, and PHE352, and polar residues CYS267 and GLN192 have closely contacted with the compound. Notably the TRP174 has formed a π-π interaction with the cyclohexane benzene like ring of the compound.

In summary, these six active compounds have similar binding style with the binding site, and the interactions are mostly dominated by the hydrophobic interaction, especially the π-related interactions. The most frequently involved interacting residues, TRP174 and TRP4, reveal their importance upon interactions.

## Discussion

### Comparing the performance of DeepGPCR_BC with other deep learning-based protein-ligand binary prediction methods

To compare the DeepGPCR_BC with other generalized protein-ligand binary prediction methods, we compared the performance of DFCNN ^12, 19, 36^ and DeepBindGCN_BC ^26^ on the 16 extra test cases. Although previously we have reported DFCNN has good performance in drug screening tasks for many solvable proteins, we found that it has notably poor performance in most of those 16 GPCR cases, listed in **Table S9**. In analogue to DeepBindGCN_BC, which shares a similar model architecture but is trained on the PDBbind database primarily consisting of soluble proteins, also showed poor performance in these 16 GPCR cases, listed in **Table S10**. The underwhelming performance of DFCNN and DeepBindGCN_BC strongly underscores the critical importance of training models like DeepGPCR_BC on GPCR ligand pairs for creating effective GPCR-ligand prediction models.

In parallel, we compared DeepGPCR_RG with the previously DeepBindGCN_RG, a graph convolutional network model trained on the PDBbind database. In order to compare it with the specialized DeepGCN_RG model developed for GPCR, we tested its performance on additional data (**Table S11**). As anticipated, DeepBindGCN_RG, having been trained on a large number of water-soluble proteins and few GPCRs, performed significantly inferior performance compared to DeepGPCR_RG on GPCR targets (**Table S11**) within the extra test set. set.

### Problem encountered when using DeepGPCR_BC and Schrödinger

Using DeepGPCR_BC and Schrödinger for small molecule screening offers certain advantages, a major limitation arises due to the extensive list of potential candidates. When thousands of predicted values are close to 1, it becomes difficult to select candidate molecules within a smaller range. Although clustering might be attempted to address this issue, the class centers may not fully capture the diverse attributes of other molecules within the same class, which could lead to the loss of effective molecules. To show this issue, we performed a small molecule screening of three proteins (O14626, O95800, and Q9HC97) using only DeepGPCR_BC and Schrödinger. We found that relying solely on DeepGPCR_BC and Schrödinger for small molecule screening results in a large data set, making it difficult to identify candidate molecules within a narrow range. Therefore, in practical applications, a more refined screening method should be adopted and combined with other tools for comprehensive analysis to ensure optimal screening performance. Detailed screening processes and results are provided in **Supplementary material section 1, Table S12 and Figure S7**.

### Importance of Combining DeepGPCR_BC and DeepGPCR_RG

As described in the result section, in order to further investigate the reliability of DeepGPCR_RG and Schrödinger’s software, we conducted experimental validation on a separate list of GPR35 candidate using DeepGPCR_RG and Schrödinger (DeepGPCR_RG > 10, Schrödinger score ≥ -6.35 kcal/mol) (**Table S7**). Remarkably, none of the 6 compounds screened illustrated any activity, depicted in **Figure 5**. This outcome strongly underscores the significance of integrating both DeepGPCR_BC and DeepGPCR_RG for effective screening of GPR35. Therefore, our findings reinforce the necessity of a comprehensive approach that synergizes multiple tools and methodologies in the process of compound screening, significantly enhancing the likelihood of identifying active compounds.

### Evaluating the reliability of Alphafold2 predicted GPCR structure

Since the GPCRs of the 16 extra test cases were all modeled by Alphafold2, it is necessary to check the accuracy of the predictions of Alphafold2 relative to the correct GPCR structure, especially the reliability of the pocket region. Here we selected 62 cases with known PDB structures and share a sequence identity larger than 0.83 with our target sequence. We used the TMalign tool ^37^ to evaluate the aligned RMSD between the Alphafold2 predicted structures and the PDB database counterparts (**Table S13**). The results indicated that most GPCRs align closely with experimental structures, exhibiting a low average RMSD of 2.19 Å, which suggests a high degree of accuracy in Alphafold2’s predictions.

### The Potential Role of MD Simulations in Analyzing Interactions

MD simulation for the GPCR with membrane lipids and ligands are relative complicated and not suitable for the large-scale screening. However, they can be particularly valuable in late-stage screening or analysis. To explore the accurate interaction details of GPR35 with the three known active compounds, (F545-1970, K297-0698, and S948-0241), we carried MD simulations and metadynamics. The comprehensive simulation procedures are outlined in **Supplementary Section 2**. We observed a consistently stable binding during the MD simulations. The analysis result of MD and metadynamics simulation for GPR35 binding with the three compounds are shown in Figure S8. The calculated RMSD values along the 40ns simulation time are relatively small, stable at around 0.2 nm, for K297-0698 and S948-0241, shown in **Figure S8B**, indicating quite stable binding. There are slight larger RMSD fluctuations for F545-1970, around 0.2∼0.6nm, but compensated with a higher number of hydrogen bonds formed during the 40ns simulation time, shown in **Figure S8B**. The calculated binding free energy landscape by metadynamics further supports the propensity of these three compounds to bind (Figure **S8C**). However, the simulation duration was relative short, and techniques like funnel metadynamcis may be more suitable for free energy calculations. It is worth noting that an in-depth exploration of MD-related methods was beyond the scope of this study.

### Improvement in future

The DeepGPCR_BC and DeepGPCR_RG models for predicting GPCR-ligand interactions can be optimized and improved in several ways. One possible way is to provide more training data, as many protein-ligand pairs were not participated to be the training data due to the absence of known ligand-binding pockets. Developing methods with high accuracy and efficiency in identifying GPCR pockets would be crucial for the inclusion of these data, thus, it can significantly expand the training set. The improved GPCR pocket identification would also enhance the model’s accuracy in screening tasks, particularly for GPCR targets without known pockets. Also, increasing the diversities of representations as input to form a multimodal may also be helpful. Optimizing hyperparameters slightly improves the capability of the model, the overestimated performance must be considered in the test set only. More sophisticated model architecture improves the performance, such as adding an attention layer, investigated by others by testing the new models in ligand property prediction ^38^.

In the screening process, the Schrödinger scores were calculated in GPR35 screening. Results in **Table 3** strongly indicating its performance of the GPCR-ligand prediction task is highly inaccurate. Therefore, it should be noted that the Schrödinger score has less reference value on the prediction, but valuable in generating GPRC-ligand complexes for MD simulations, and a more accurate affinity prediction method should be developed. Additionally, a more comprehensive pipeline could be constructed to categorize high-potential candidates step-by-step, for examples, integrating methods such as molecular dynamics (MD) simulations and metadynamics to identify reliable candidates. Finally, experimental validations will be essential in assessing the usefulness of such predictions.

Deep-learning-based methods usually demonstrating the black-box property, in fact, considering the interpretability of the DeepGPCR_BC and DeepGPCR_RG models in specific problem-solving is extremely important and guide to the potential drug candidate optimization. Here, we recommend two possible solutions to relieve the black-box property of DeepGPCR_BC and DeepGPCR_RG. First, the GCN model has an advantage in interpreting each node’s contributions to the prediction. In other words, the atoms in ligands or pocket residues contributions can be revealed, then the hot spot atom or residue can be detected. Users are able to achieve and visualize the atom contributions by RdKit tools. Researchers have provided several methods ^39^ with relevant scripts (https://github.com/biomed-AI/MolRep). The second way is to combine the DeepGPCR_BC and DeepGPCR_RG with the docking tools or MD simulation tools to further explore the atomic binding details and compare the binding pose with the known GPCR-drug complexes. Employing MD simulations to ensure the binding stability and applying Funnel metadynamics ^40^ methods to calculate the binding free energy of interested protein-ligand pairs obtained by DeepGPCR_BC and DeepGPCR_RG are recommended. But such a method is relatively time and resource-consuming, and GPCR simulation needs to incorporate a large number of lipids and solvents, the simulations are especially complicated.

## Conclusion

The G protein-coupled receptor (GPCR) is a critical drug target, the traditional protein-ligand interaction prediction software, however, are unsuitable for GPCR drug virtual screening due to its unique properties and environment. To narrow this gap, a specific GPCR-ligand interaction model has been developed in this work by learning the underlying interacting rules between the GPCR-ligand system, with inputting the GPCR pocket and ligand separately. The model employs graph representations denoting the protein pocket and ligand, with residues or atoms as nodes and contacting residues or bonds as edges. Each pocket and ligand input are submitted into the graphic neural network, and the final output is merged. The model has been trained on a huge GPCR dataset with no available GPCR-ligand complex structures. This advantage is significant over traditional docking or generalized models trained on mostly non-GPCR-ligand datasets. This model fully utilizes the spatial and physical-chemical features of GPCR pockets and known ligands, making it more suitable for screening active compounds for GPCR. The DeepGPCR_BC model has achieved an average of 0.72, 0.73, and 0.73 for AUC, TPR (recall), and precision, respectively. In the 16 fully independent test sets, our model exhibits a significantly superior performance compared to the scores from Schrödinger and Autodock Vina, with a cutoff of -6 kcal/mol. Notably, 9 out of the 16 cases achieved an AUC greater than 0.7, and 4 cases achieved an AUC above 0.8. It should be noted that a few cases have not performed well, partly due to the challenges of pocket identification. Also, the performance of our model can be further enhanced when more GPCRs with identified ligand pockets are available. The DeepGPCR_RG model has achieved an average 0.39 Pearson correlation, much better performance than Schrödinger and Autodock Vina, also superior than our previous DeepBindGCN_RG, in the 16-protein related extra GPCR test dataset. Additionally, the development of an affinity prediction model would help narrow the candidate list and aid in obtaining more reliable and high-affinity candidates. Most importantly, we have built a screening pipeline use DeepGPCR models as core components, and successfully applied to two important GPCR therapeutic target GPR35 and GLP_R1, resulting 3 active compounds out of 8 candidates for GPR35 by strategy A, 5 active compounds out of 12 candidates for GLP_R1 by strategy A, and 1 out of 3 candidates by Strategy B. Overall, the DeepGPCR_BC and DeepGPCR_RG models provide promising advancement in the field of GPCR drug discovery, facilitating the identification of novel GPCR drugs and enhancing the precision of GPCR drug virtual screening.

## Data Availability Statement

The proposed DeepGPCR_BC model and the scripts are available in GitHub public repositories (https://github.com/haiping1010/DeepGPCR). The proposed DeepGPCR_RG model and its accompanying scripts are available upon appropriate request through the corresponding author.

## Author contributions

HZ and YC designed the study. HZ performed the coding, model construction, and computations. HF, JW and TH performed the experiments, HZ, YC, JW, TH, HF, KMS, JL, WX and KHW performed the data analysis. HZ, YC, XL and JZ supervised the study. HZ and YC wrote the manuscript with input from all other authors. All authors read and approved the final manuscript.

## Acknowledgments

This study was supported by the National Science Foundation of China (62106253, 21933010, 32060209, 82103187, 22250710136), Shenzhen Key Projects (JCYJ20220818100804009), Jiangxi Provincial Natural Science Foundation of China (20232BBH80012, 20224ACB216012), Liaoning Provincial Natural Science Foundation of China (2022-MS-017) and Dalian High-Level Talent Support Program (2021RQ003).

## Competing Interests

No authors have a conflict of interest in publishing this paper.

## Supplementary Figures

**Figure S1.**
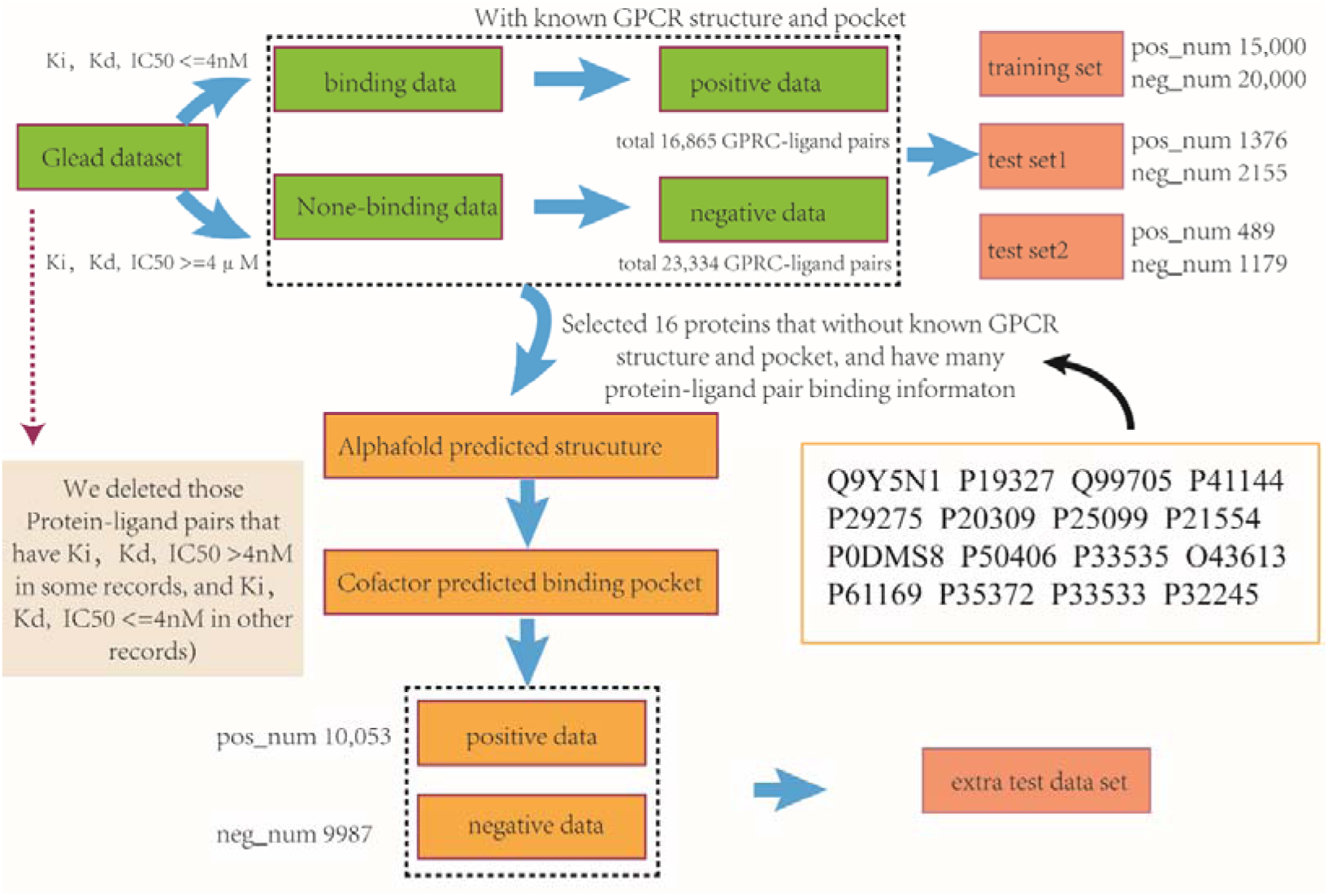
Data preparation for training and test.

**Figure S2.**
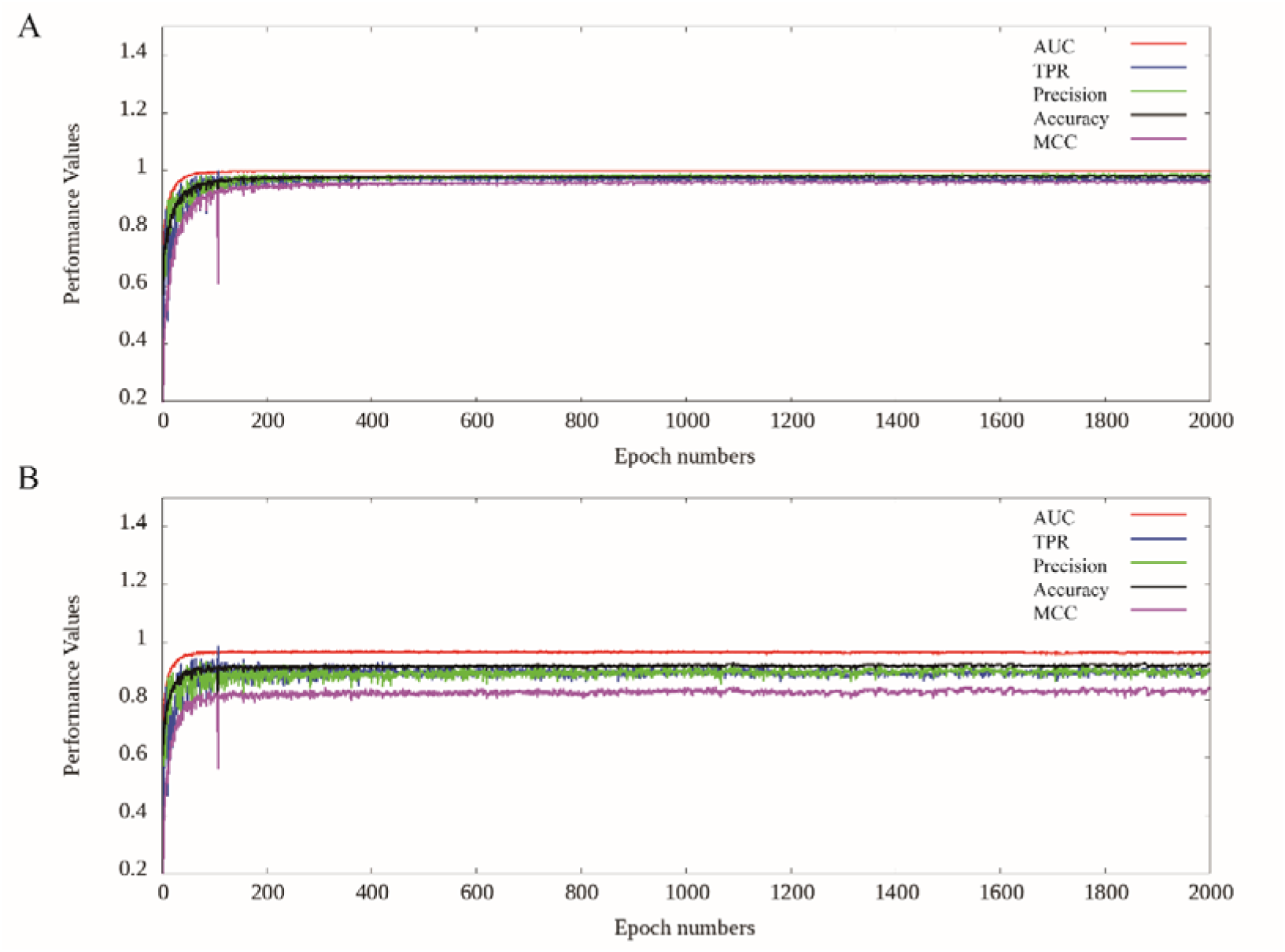
The plot of model performance on the training and testing set along different training epochs.

**Figure S3.**
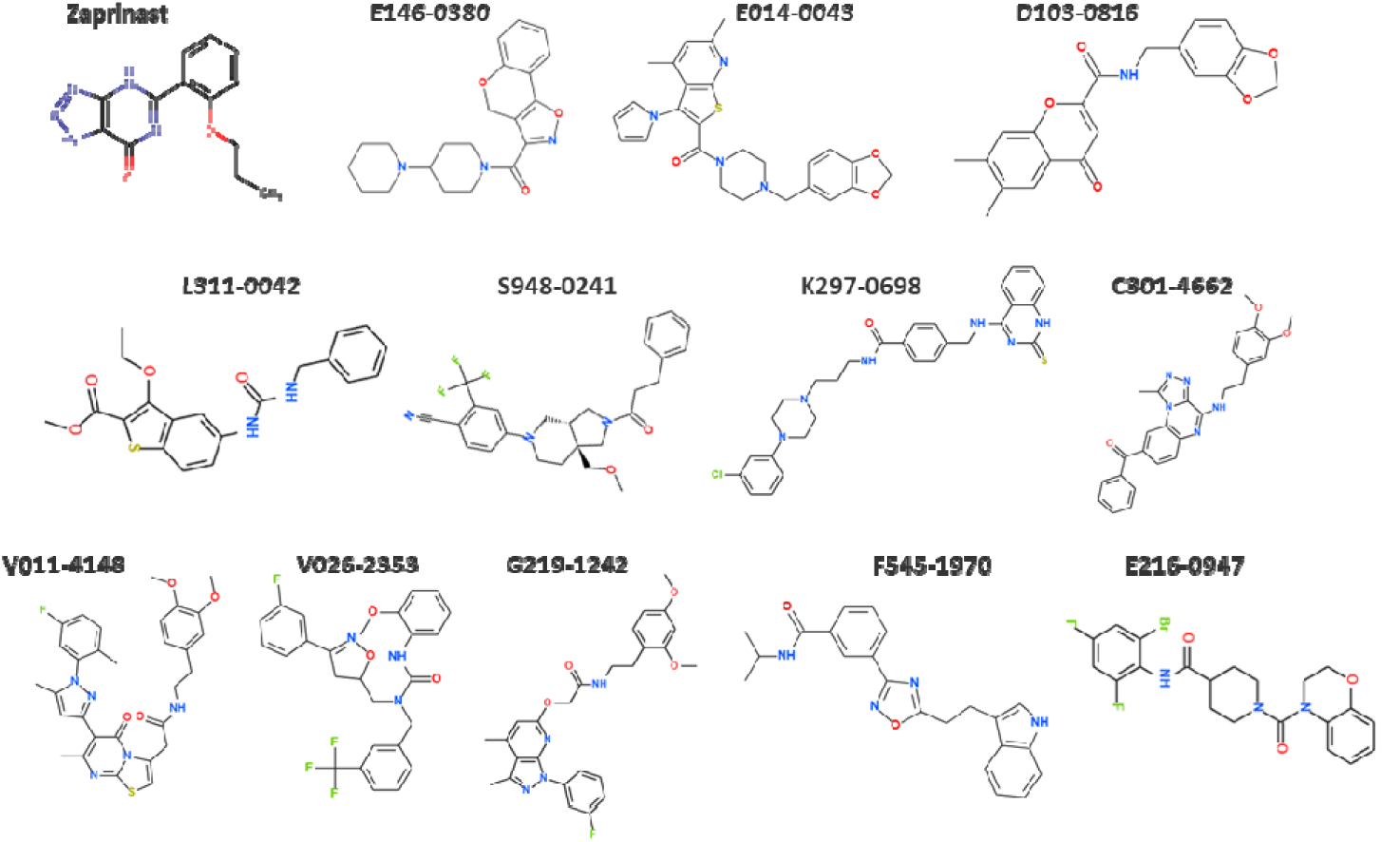
Chemical structures of 12 selective candidates for GPR35.

**Figure S4.**
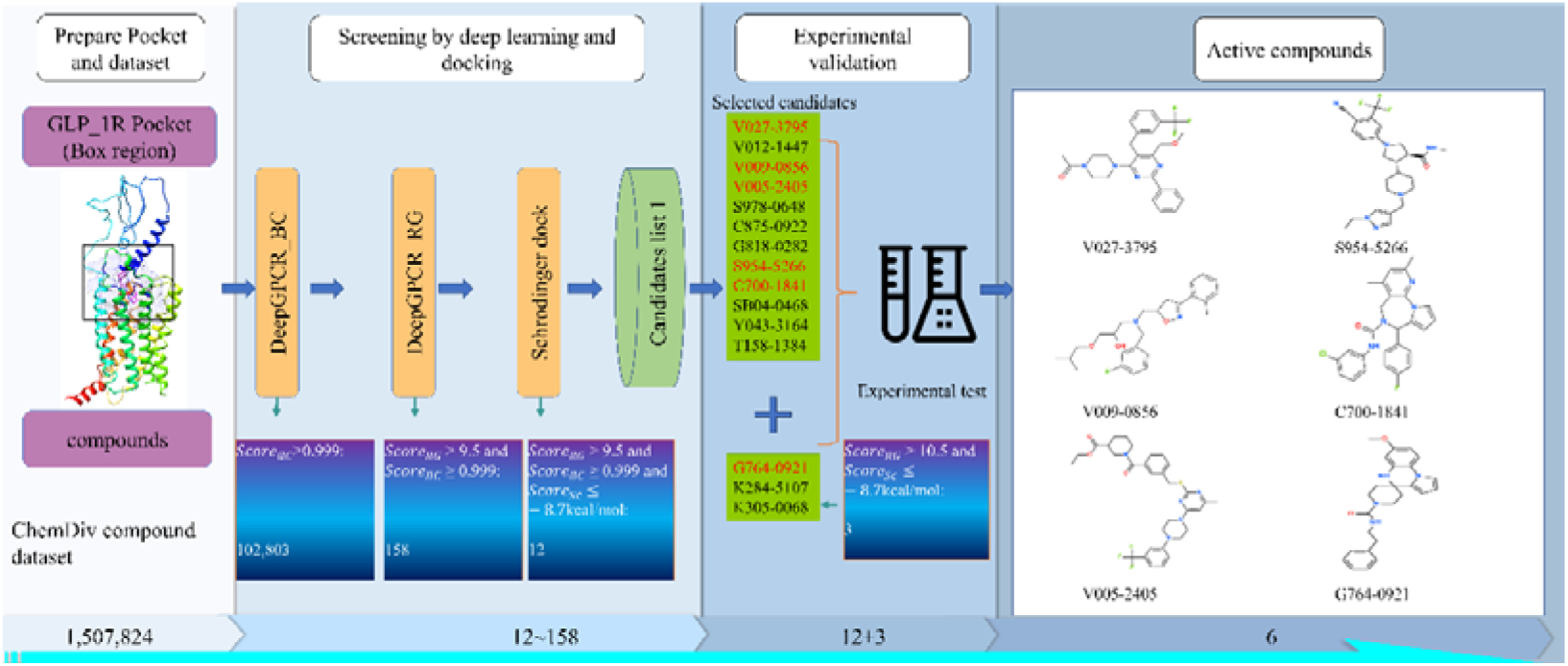
GLP_1R screening pipeline and Identification of Active Compounds. Schematic representation of the stepwise screening process leading to the discovery of 6 active molecules.

**Figure S5.**
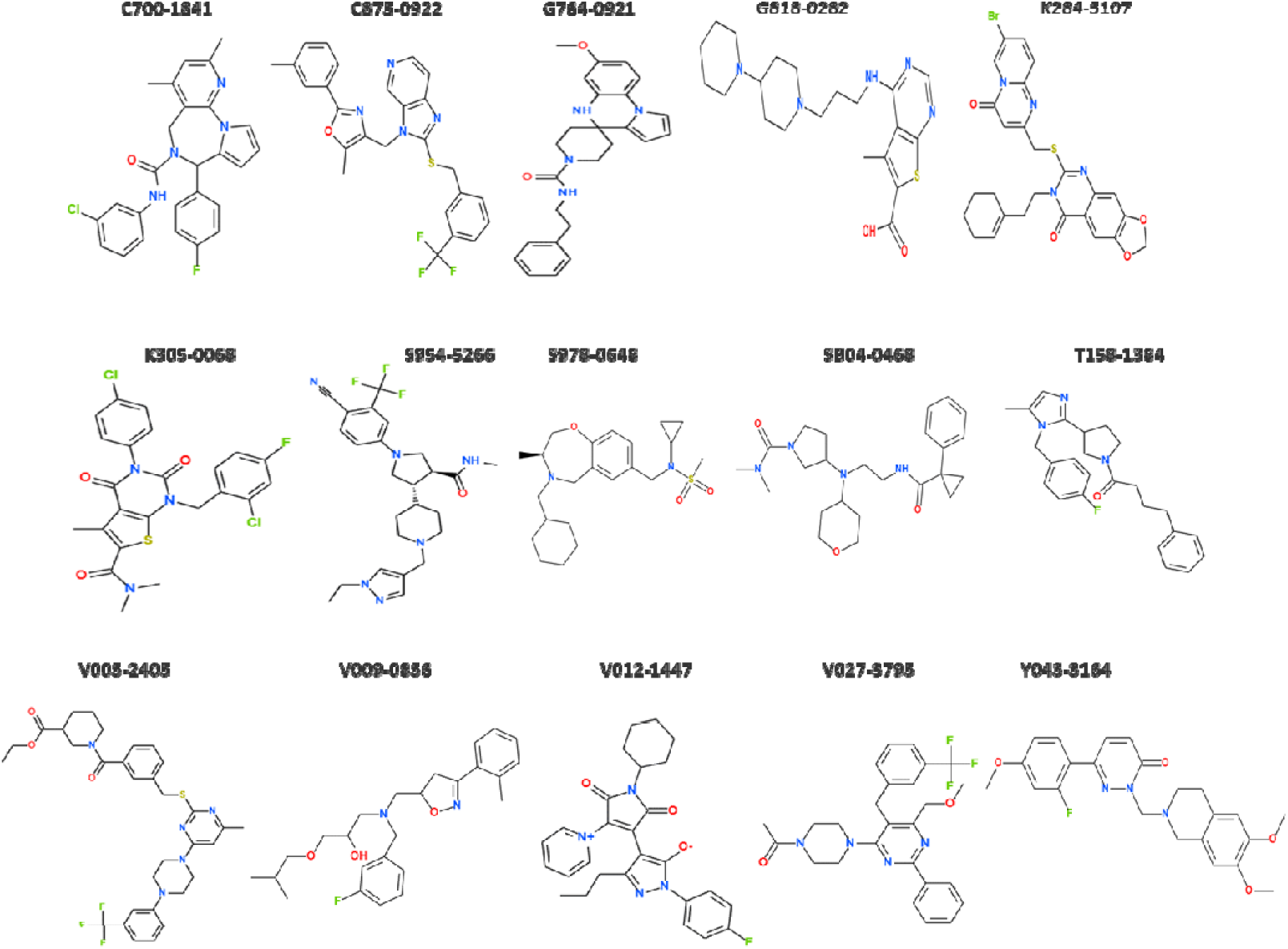
Chemical structures of 15 selective candidates for GLP-1R.

**Figure S6.**
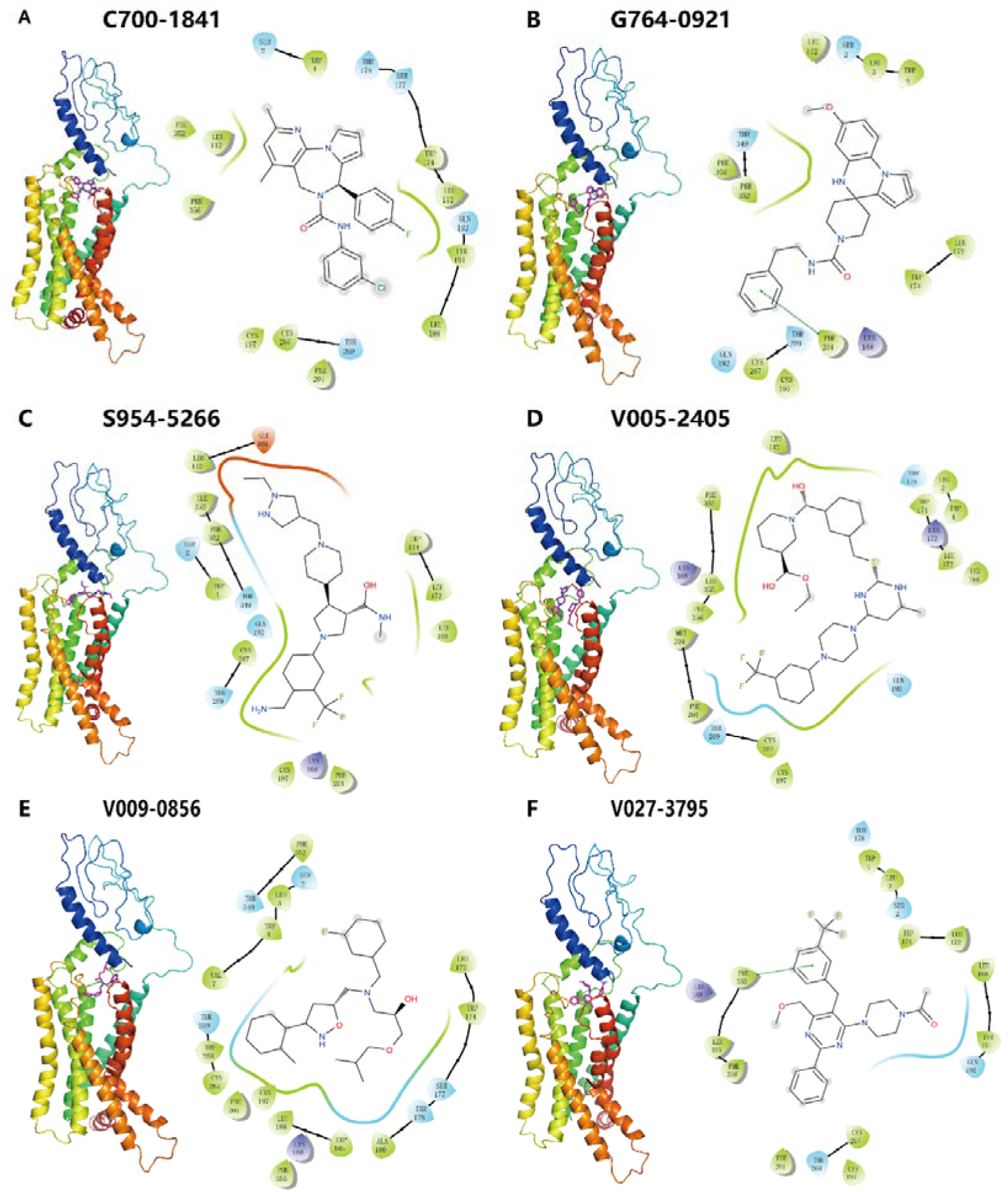
the overall binding view and 2D plot of GLP_R1 with six active molecules from docking. A, interactions between pocket residues and compound C700-1841. B. interactions between pocket residues and compound G764-0921. C. interactions between pocket residues and compound S954-5266. D. interactions between pocket residues and known active compound V005-2405. E. interactions between pocket residues and known active compound V009-0856. F. interactions between pocket residues and known active compound V027-3795.

**Figure S7.**
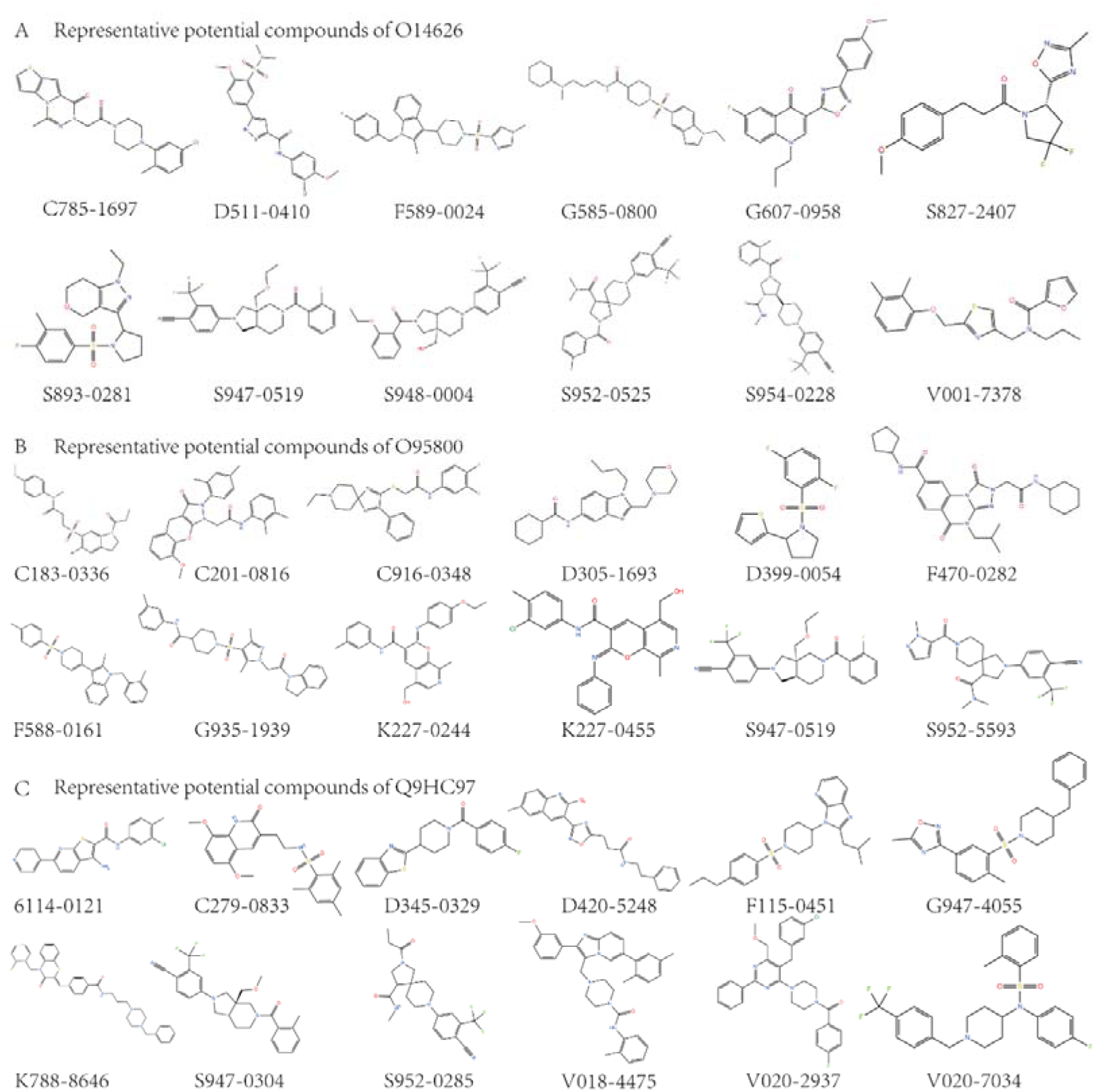
The potential representative compounds of O14626, O95800, and Q9HC97, respectively. A. Many of the representative compound candidates of O14626 show a linear shape. B. The structure of the representative compound candidates of O958000 is relatively diversified. C. The representative compounds candidates of Q9HC97, several representative structures contain common chemical groups, such as sulfonyl.

**Figure S8.**
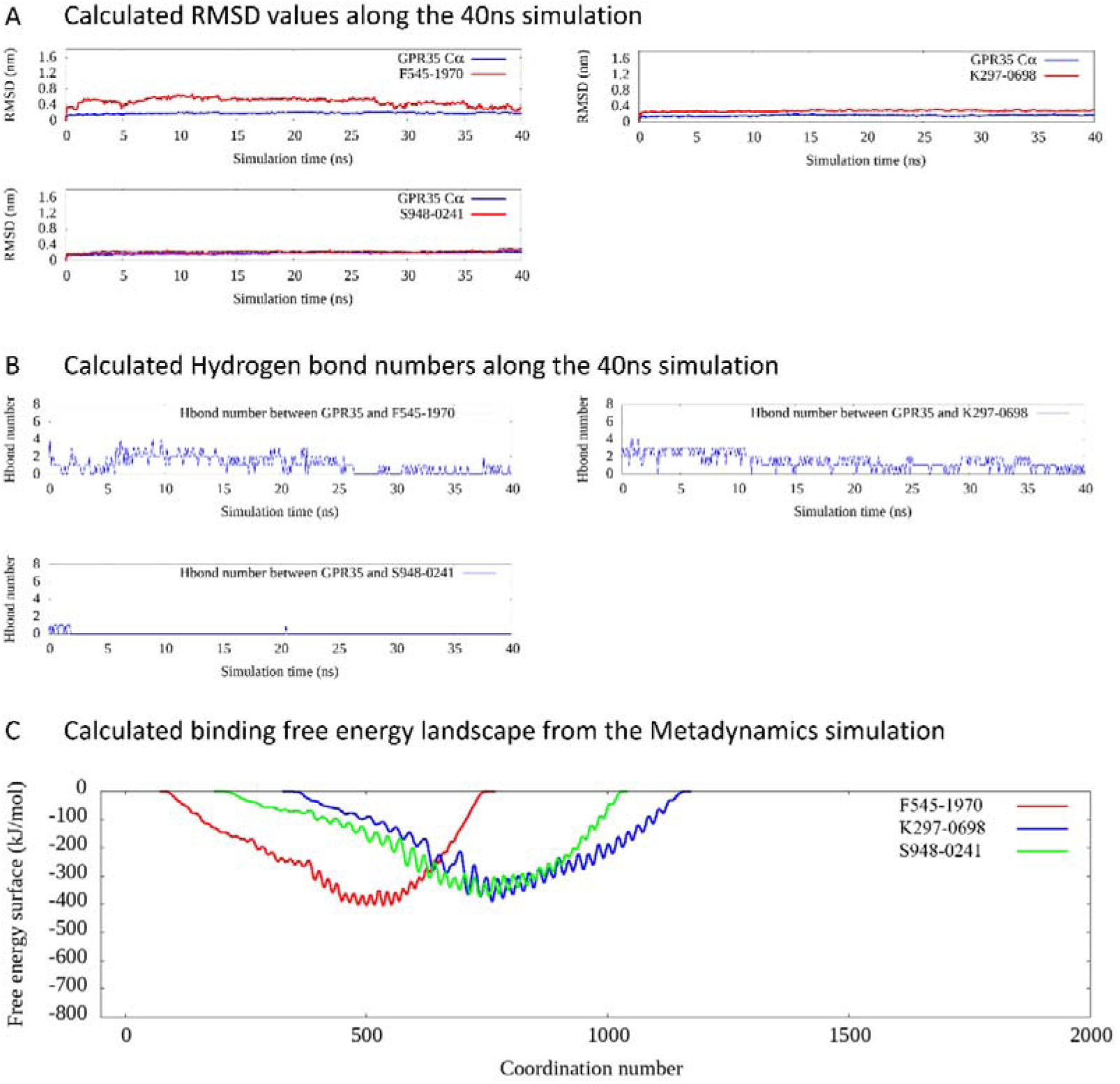
The analysis result of MD and metadynamics simulation for GPR35 binding with the three compounds (F545-1970 K297-0698 and S948-0241). A. The calculated RMSD value along the 40ns simulation time; B. The calculated hydrogen bond number along the 40ns simulation time; C. The calculated binding free energy landscape by metadynamics.

## Supplementary Tables

**Table S1.** The model performance on the training and test set1 at different training epochs, measured by AUC, TPR, precision, accuracy, and MCC.

| Dataset | epoch | AUC | TPR | Precision | Accuracy | MCC |
| --- | --- | --- | --- | --- | --- | --- |
| Training set | 100 | 1.00 | 0.97 | 0.95 | 0.97 | 0.93 |
|  | 200 | 1.00 | 0.97 | 0.96 | 0.97 | 0.94 |
|  | 300 | 1.00 | 0.97 | 0.97 | 0.98 | 0.95 |
|  | 400 | 1.00 | 0.98 | 0.97 | 0.98 | 0.95 |
|  | 500 | 1.00 | 0.98 | 0.98 | 0.98 | 0.96 |
|  | 600 | 1.00 | 0.97 | 0.98 | 0.98 | 0.96 |
|  | 700 | 1.00 | 0.97 | 0.98 | 0.98 | 0.96 |
|  | 800 | 1.00 | 0.97 | 0.98 | 0.98 | 0.96 |
|  | 900 | 1.00 | 0.97 | 0.97 | 0.98 | 0.95 |
|  | 1000 | 1.00 | 0.97 | 0.99 | 0.98 | 0.96 |
|  | 1100 | 1.00 | 0.97 | 0.99 | 0.98 | 0.96 |
|  | 1200 | 1.00 | 0.97 | 0.98 | 0.98 | 0.96 |
|  | 1300 | 1.00 | 0.97 | 0.98 | 0.98 | 0.96 |
|  | 1400 | 1.00 | 0.97 | 0.98 | 0.98 | 0.96 |
|  | 1500 | 1.00 | 0.97 | 0.99 | 0.98 | 0.96 |
|  | 1600 | 1.00 | 0.97 | 0.99 | 0.98 | 0.96 |
|  | 1700 | 1.00 | 0.97 | 0.99 | 0.98 | 0.96 |
|  | 1800 | 1.00 | 0.97 | 0.99 | 0.98 | 0.96 |
|  | 1900 | 1.00 | 0.97 | 0.99 | 0.98 | 0.96 |
|  | 2000 | 1.00 | 0.98 | 0.98 | 0.98 | 0.96 |
| Test set1 | 100 | 0.96 | 0.92 | 0.87 | 0.91 | 0.82 |
|  | 200 | 0.97 | 0.90 | 0.87 | 0.91 | 0.81 |
|  | 300 | 0.97 | 0.90 | 0.89 | 0.92 | 0.82 |
|  | 400 | 0.97 | 0.92 | 0.86 | 0.91 | 0.82 |
|  | 500 | 0.97 | 0.90 | 0.89 | 0.92 | 0.83 |
|  | 600 | 0.97 | 0.90 | 0.88 | 0.91 | 0.82 |
|  | 700 | 0.97 | 0.90 | 0.89 | 0.92 | 0.83 |
|  | 800 | 0.97 | 0.88 | 0.89 | 0.91 | 0.82 |
|  | 900 | 0.97 | 0.91 | 0.87 | 0.92 | 0.82 |
|  | 1000 | 0.97 | 0.90 | 0.90 | 0.92 | 0.83 |
|  | 1100 | 0.97 | 0.89 | 0.91 | 0.92 | 0.84 |
|  | 1200 | 0.97 | 0.89 | 0.89 | 0.92 | 0.82 |
|  | 1300 | 0.96 | 0.90 | 0.88 | 0.91 | 0.82 |
|  | 1400 | 0.96 | 0.90 | 0.89 | 0.92 | 0.82 |
|  | 1500 | 0.97 | 0.90 | 0.90 | 0.92 | 0.84 |
|  | 1600 | 0.97 | 0.90 | 0.90 | 0.92 | 0.84 |
|  | 1700 | 0.96 | 0.89 | 0.90 | 0.92 | 0.83 |
|  | 1800 | 0.96 | 0.89 | 0.90 | 0.92 | 0.83 |
|  | 1900 | 0.96 | 0.90 | 0.90 | 0.92 | 0.83 |
|  | 2000 | 0.97 | 0.90 | 0.90 | 0.92 | 0.84 |

**Table S2.** The performance on the test set2 using model at 2000th training epoch, measured by AUC, TPR, precision, accuracy, and MCC. Notably, this set only contain protein P29274 related interaction.

| AUC | TPR | Precision | Accuracy | MCC | pos_num | neg_num |
| --- | --- | --- | --- | --- | --- | --- |
| 0.72 | 0.46 | 0.46 | 0.70 | 0.16 | 489 | 1179 |

**Table S3.**
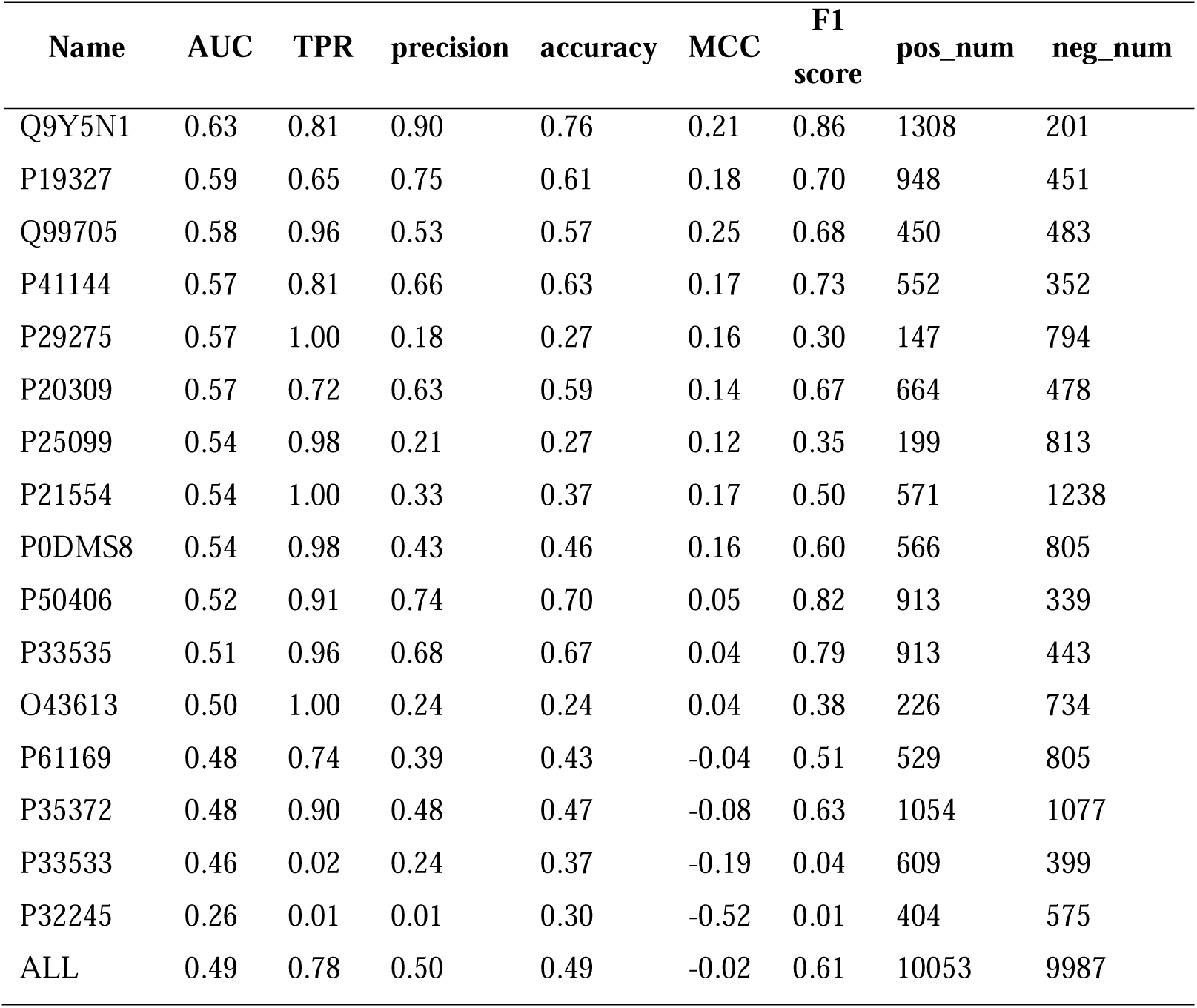
The Autodock vina performance on an extra dataset with modeled GPCR protein and predicted pocket. We used -6 Kcal/mol as the cutoff, those scores > -6 Kcal/mol was assigned a value of 0 (indicating non-bind), and those scores ≤ -6 Kcal/mol was assigned a value of 1 (indicating able to bind).

| Name | AUC | TPR | precision | accuracy | MCC | F1 score | pos_num | neg_num |
| --- | --- | --- | --- | --- | --- | --- | --- | --- |
| Q9Y5N1 | 0.63 | 0.81 | 0.90 | 0.76 | 0.21 | 0.86 | 1308 | 201 |
| P19327 | 0.59 | 0.65 | 0.75 | 0.61 | 0.18 | 0.70 | 948 | 451 |
| Q99705 | 0.58 | 0.96 | 0.53 | 0.57 | 0.25 | 0.68 | 450 | 483 |
| P41144 | 0.57 | 0.81 | 0.66 | 0.63 | 0.17 | 0.73 | 552 | 352 |
| P29275 | 0.57 | 1.00 | 0.18 | 0.27 | 0.16 | 0.30 | 147 | 794 |
| P20309 | 0.57 | 0.72 | 0.63 | 0.59 | 0.14 | 0.67 | 664 | 478 |
| P25099 | 0.54 | 0.98 | 0.21 | 0.27 | 0.12 | 0.35 | 199 | 813 |
| P21554 | 0.54 | 1.00 | 0.33 | 0.37 | 0.17 | 0.50 | 571 | 1238 |
| P0DMS8 | 0.54 | 0.98 | 0.43 | 0.46 | 0.16 | 0.60 | 566 | 805 |
| P50406 | 0.52 | 0.91 | 0.74 | 0.70 | 0.05 | 0.82 | 913 | 339 |
| P33535 | 0.51 | 0.96 | 0.68 | 0.67 | 0.04 | 0.79 | 913 | 443 |
| O43613 | 0.50 | 1.00 | 0.24 | 0.24 | 0.04 | 0.38 | 226 | 734 |
| P61169 | 0.48 | 0.74 | 0.39 | 0.43 | -0.04 | 0.51 | 529 | 805 |
| P35372 | 0.48 | 0.90 | 0.48 | 0.47 | -0.08 | 0.63 | 1054 | 1077 |
| P33533 | 0.46 | 0.02 | 0.24 | 0.37 | -0.19 | 0.04 | 609 | 399 |
| P32245 | 0.26 | 0.01 | 0.01 | 0.30 | -0.52 | 0.01 | 404 | 575 |
| ALL | 0.49 | 0.78 | 0.50 | 0.49 | -0.02 | 0.61 | 10053 | 9987 |

**Table S4.** Performance of DeepGPCR_RG for Training and test set at different epochs.

| **Dataset** | **epoch** | **rmse** | **mse** | **pearson** | **spearman** |
| --- | --- | --- | --- | --- | --- |
| Training | 200 | 0.72 | 0.51 | 0.84 | 0.83 |
|  | 400 | 0.68 | 0.46 | 0.86 | 0.85 |
|  | 600 | 0.66 | 0.44 | 0.87 | 0.86 |
|  | 800 | 0.66 | 0.44 | 0.87 | 0.86 |
|  | 1000 | 0.66 | 0.43 | 0.87 | 0.86 |
|  | 1200 | 0.65 | 0.42 | 0.87 | 0.86 |
|  | 1400 | 0.66 | 0.43 | 0.87 | 0.86 |
|  | 1600 | 0.65 | 0.42 | 0.87 | 0.86 |
|  | 1800 | 0.65 | 0.42 | 0.87 | 0.86 |
|  | 2000 | 0.64 | 0.41 | 0.87 | 0.86 |
| test | 200 | 0.98 | 0.96 | 0.68 | 0.67 |
|  | 400 | 1.00 | 1.01 | 0.67 | 0.65 |
|  | 600 | 1.01 | 1.03 | 0.66 | 0.64 |
|  | 800 | 1.04 | 1.09 | 0.66 | 0.64 |
|  | 1000 | 1.02 | 1.04 | 0.65 | 0.63 |
|  | 1200 | 1.04 | 1.09 | 0.65 | 0.63 |
|  | 1400 | 1.03 | 1.06 | 0.64 | 0.63 |
|  | 1600 | 1.04 | 1.08 | 0.64 | 0.63 |
|  | 1800 | 1.05 | 1.11 | 0.64 | 0.63 |
|  | 2000 | 1.05 | 1.11 | 0.64 | 0.63 |

**Table S5.** The Schrödinger docking performance on regression model’s extra test datasets.

| Name | rmse | mse | pearson | spearman | total_num |
| --- | --- | --- | --- | --- | --- |
| Q9Y5N1 | 4.93 | 24.31 | 0.16 | 0.17 | 3693 |
| P61169 | 3.55 | 12.59 | 0.10 | 0.08 | 4460 |
| P29275 | 2.43 | 5.91 | 0.09 | 0.17 | 1804 |
| P19327 | 3.85 | 14.83 | 0.06 | 0.09 | 4632 |
| P33535 | 3.92 | 15.39 | 0.02 | 0.02 | 2962 |
| P25099 | 1.83 | 3.33 | 0.01 | -0.01 | 2932 |
| P21554 | 3.73 | 13.94 | 0.01 | -0.02 | 3530 |
| P0DMS8 | 2.15 | 4.60 | 0.00 | -0.02 | 3797 |
| P20309 | 4.57 | 20.89 | -0.01 | 0.01 | 2006 |
| Q99705 | 4.34 | 18.86 | -0.02 | 0.05 | 3610 |
| P50406 | 3.78 | 14.30 | -0.02 | -0.01 | 3297 |
| O43613 | 5.73 | 32.86 | -0.06 | -0.06 | 3811 |
| P41144 | 5.10 | 26.00 | -0.08 | -0.02 | 2173 |
| P35372 | 3.62 | 13.10 | -0.09 | -0.06 | 4236 |
| P33533 | 7.62 | 58.10 | -0.13 | -0.12 | 1969 |
| P32245 | 7.01 | 49.17 | -0.30 | -0.26 | 2613 |
| Average | 4.37 | 19.11 | -0.04 | -0.04 | 51525 |

**Table S6.** The vina docking performance on regression model’s extra test datasets.

| **Name** | **rmse** | **mse** | **pearson** | **spearman** | **total_num** |
| --- | --- | --- | --- | --- | --- |
| O43613 | 2.02 | 4.09 | 0.02 | 0.11 | 3811 |
| P19327 | 3.22 | 10.39 | 0.05 | 0.06 | 4632 |
| P21554 | 1.81 | 3.28 | 0.25 | 0.26 | 3530 |
| P29275 | 1.71 | 2.94 | 0.31 | 0.36 | 1804 |
| P33533 | 10.75 | 115.46 | 0.08 | -0.11 | 1969 |
| P41144 | 3.30 | 10.88 | 0.05 | 0.04 | 2173 |
| Q99705 | 2.31 | 5.32 | 0.18 | 0.18 | 3610 |
| P33535 | 3.73 | 13.91 | -0.01 | 0.05 | 2962 |
| P50406 | 2.56 | 6.57 | 0.02 | 0.02 | 3297 |
| Q9Y5N1 | 3.40 | 11.59 | 0.19 | 0.21 | 3693 |
| P0DMS8 | 2.02 | 4.10 | -0.04 | -0.08 | 3797 |
| P20309 | 4.13 | 17.02 | 0.03 | 0.10 | 2006 |
| P25099 | 1.62 | 2.64 | 0.13 | 0.13 | 2932 |
| P32245 | 15.00 | 225.02 | -0.29 | -0.42 | 2613 |
| P35372 | 4.13 | 17.09 | -0.05 | -0.03 | 4236 |
| P61169 | 2.40 | 5.77 | 0.03 | 0.08 | 4460 |
| Average | 4.83 | 23.29 | -0.06 | -0.07 | 51525 |

**Table S7.** The compound list of GPR35 by using DeepGPCR_RG and Schrödinger (DeepGPCR_RG≥10, Schrödinger score≤-6.35 Kcal/mol).

| Chemdiv id | DeepGPCR_RG | Schrödinger<br>(Kcal/mol) |
| --- | --- | --- |
| V010-1264 | 10.03 | -7.02 |
| 6028-3969 | 10.30 | -6.77 |
| E146-0380 | 10.45 | -6.47 |
| E014-0043 | 12.16 | -6.40 |
| D103-0816 | 10.10 | -6.39 |
| L311-0042 | 10.18 | -6.35 |

**Table S8.** The compound list of GLP_1R by using DeepGPCR_RG and Schrödinger (DeepGPCR_RG≥10.5, Schrödinger score≤-8.7 Kcal/mol).

| Chemdiv id | DeepGPCR_RG | Schrödinger<br>(Kcal/mol) |
| --- | --- | --- |
| G764-0921 | 10.80 | -8.71 |
| K284-5107 | 10.71 | -8.90 |
| K305-0068 | 10.61 | -8.79 |

**Table S9.** DFCNN performance on an extra dataset with modeled GPCR protein and predicted pocket.

| Name | AUC | TPR | precision | accuracy | MCC | F1<br>score | pos_num | neg_num |
| --- | --- | --- | --- | --- | --- | --- | --- | --- |
| P29275 | 0.75 | 0.56 | 0.47 | 0.83 | 0.42 | 0.52 | 147 | 794 |
| P35372 | 0.66 | 0.42 | 0.65 | 0.61 | 0.22 | 0.51 | 1054 | 1105 |
| P21554 | 0.63 | 0.02 | 0.09 | 0.63 | -0.13 | 0.03 | 571 | 1242 |
| P25099 | 0.60 | 0.25 | 0.20 | 0.66 | 0.01 | 0.22 | 199 | 833 |
| P0DMS8 | 0.57 | 0.14 | 0.32 | 0.53 | -0.08 | 0.19 | 566 | 813 |
| P33533 | 0.57 | 0.63 | 0.63 | 0.56 | 0.09 | 0.63 | 609 | 409 |
| P61169 | 0.55 | 0.61 | 0.43 | 0.53 | 0.09 | 0.51 | 529 | 817 |
| Q9Y5N1 | 0.51 | 0.29 | 0.82 | 0.34 | -0.07 | 0.43 | 1308 | 219 |
| P32245 | 0.50 | 0.38 | 0.43 | 0.53 | 0.02 | 0.40 | 404 | 575 |
| P33535 | 0.49 | 0.33 | 0.61 | 0.41 | -0.10 | 0.43 | 913 | 449 |
| P50406 | 0.48 | 0.25 | 0.73 | 0.38 | 0.00 | 0.37 | 913 | 339 |
| Q99705 | 0.43 | 0.76 | 0.46 | 0.46 | -0.07 | 0.58 | 450 | 485 |
| P19327 | 0.37 | 0.19 | 0.50 | 0.32 | -0.23 | 0.28 | 948 | 453 |
| P41144 | 0.33 | 0.06 | 0.49 | 0.39 | -0.07 | 0.11 | 552 | 362 |
| O43613 | 0.23 | 0.10 | 0.07 | 0.45 | -0.30 | 0.08 | 226 | 734 |
| P20309 | 0.14 | 0.10 | 0.19 | 0.24 | -0.52 | 0.13 | 664 | 492 |
| ALL | 0.55 | 0.31 | 0.49 | 0.49 | -0.01 | 0.38 | 10053 | 10121 |

**Table S10.**
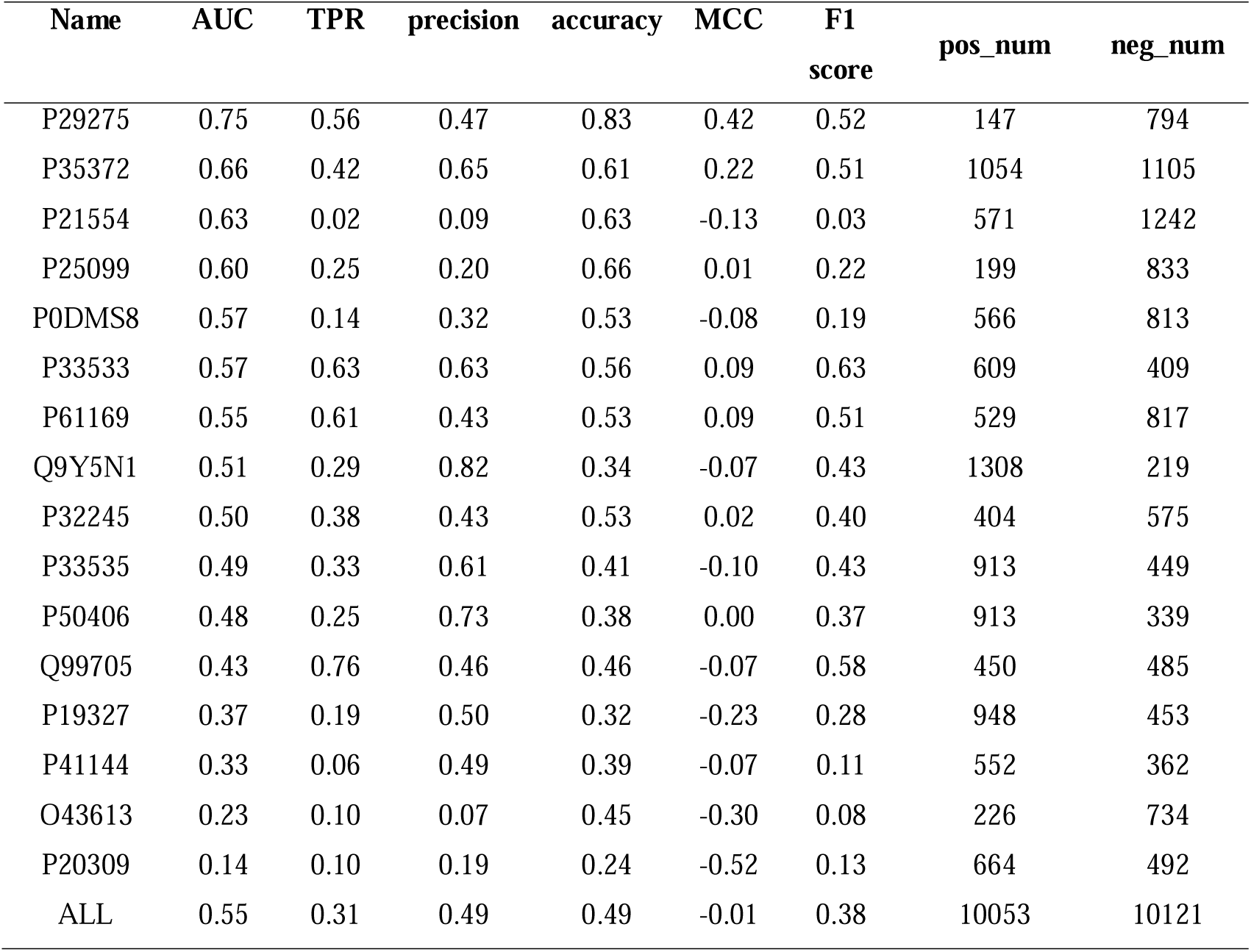
DeepBindGCN_BC performance on an extra dataset with modeled GPCR protein and predicted pocket.

| Name | AUC | TPR | precision | accuracy | MCC | F1<br>score | pos_num | neg_num |
| --- | --- | --- | --- | --- | --- | --- | --- | --- |
| P29275 | 0.75 | 0.56 | 0.47 | 0.83 | 0.42 | 0.52 | 147 | 794 |
| P35372 | 0.66 | 0.42 | 0.65 | 0.61 | 0.22 | 0.51 | 1054 | 1105 |
| P21554 | 0.63 | 0.02 | 0.09 | 0.63 | -0.13 | 0.03 | 571 | 1242 |
| P25099 | 0.60 | 0.25 | 0.20 | 0.66 | 0.01 | 0.22 | 199 | 833 |
| P0DMS8 | 0.57 | 0.14 | 0.32 | 0.53 | -0.08 | 0.19 | 566 | 813 |
| P33533 | 0.57 | 0.63 | 0.63 | 0.56 | 0.09 | 0.63 | 609 | 409 |
| P61169 | 0.55 | 0.61 | 0.43 | 0.53 | 0.09 | 0.51 | 529 | 817 |
| Q9Y5N1 | 0.51 | 0.29 | 0.82 | 0.34 | -0.07 | 0.43 | 1308 | 219 |
| P32245 | 0.50 | 0.38 | 0.43 | 0.53 | 0.02 | 0.40 | 404 | 575 |
| P33535 | 0.49 | 0.33 | 0.61 | 0.41 | -0.10 | 0.43 | 913 | 449 |
| P50406 | 0.48 | 0.25 | 0.73 | 0.38 | 0.00 | 0.37 | 913 | 339 |
| Q99705 | 0.43 | 0.76 | 0.46 | 0.46 | -0.07 | 0.58 | 450 | 485 |
| P19327 | 0.37 | 0.19 | 0.50 | 0.32 | -0.23 | 0.28 | 948 | 453 |
| P41144 | 0.33 | 0.06 | 0.49 | 0.39 | -0.07 | 0.11 | 552 | 362 |
| O43613 | 0.23 | 0.10 | 0.07 | 0.45 | -0.30 | 0.08 | 226 | 734 |
| P20309 | 0.14 | 0.10 | 0.19 | 0.24 | -0.52 | 0.13 | 664 | 492 |
| ALL | 0.55 | 0.31 | 0.49 | 0.49 | -0.01 | 0.38 | 10053 | 10121 |

**Table S11.** DeepBindGCN_RG performance on the 16-protein related extra dataset.

| **Name** | **rmse** | **mse** | **pearson** | **spearman** | **total_num** |
| --- | --- | --- | --- | --- | --- |
| P29275 | 1.27 | 1.62 | 0.31 | 0.29 | 1804 |
| Q9Y5N1 | 1.30 | 1.69 | 0.10 | 0.12 | 3693 |
| P50406 | 1.45 | 2.11 | 0.11 | 0.10 | 3297 |
| P25099 | 1.67 | 2.80 | 0.08 | 0.09 | 2932 |
| P35372 | 1.66 | 2.75 | 0.27 | 0.27 | 4236 |
| O43613 | 1.16 | 1.35 | 0.20 | 0.20 | 3811 |
| P0DMS8 | 1.45 | 2.09 | 0.21 | 0.20 | 3797 |
| P21554 | 1.15 | 1.33 | 0.31 | 0.29 | 3530 |
| P33533 | 1.54 | 2.37 | 0.08 | 0.07 | 1969 |
| P41144 | 1.30 | 1.70 | 0.28 | 0.28 | 2173 |
| P32245 | 1.52 | 2.31 | 0.22 | 0.20 | 2613 |
| P20309 | 1.68 | 2.83 | 0.28 | 0.29 | 2006 |
| Q99705 | 1.25 | 1.56 | 0.22 | 0.22 | 3610 |
| P33535 | 1.54 | 2.37 | 0.20 | 0.21 | 2962 |
| P61169 | 1.32 | 1.75 | -0.01 | -0.02 | 4460 |
| P19327 | 1.63 | 2.65 | 0.07 | 0.06 | 4632 |
| Average | 1.44 | 2.07 | 0.20 | 0.19 | 51525 |

**Table S12.**
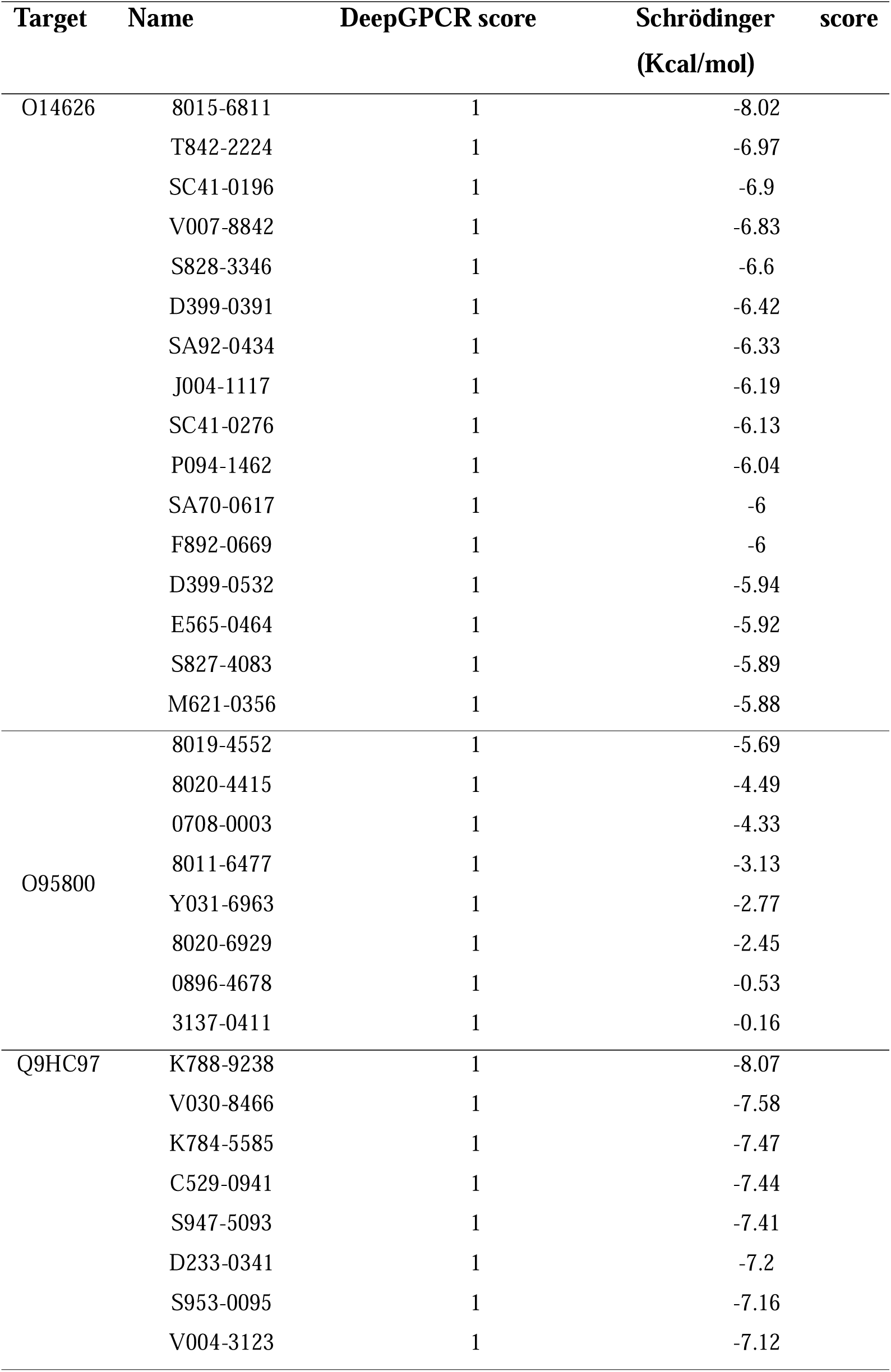
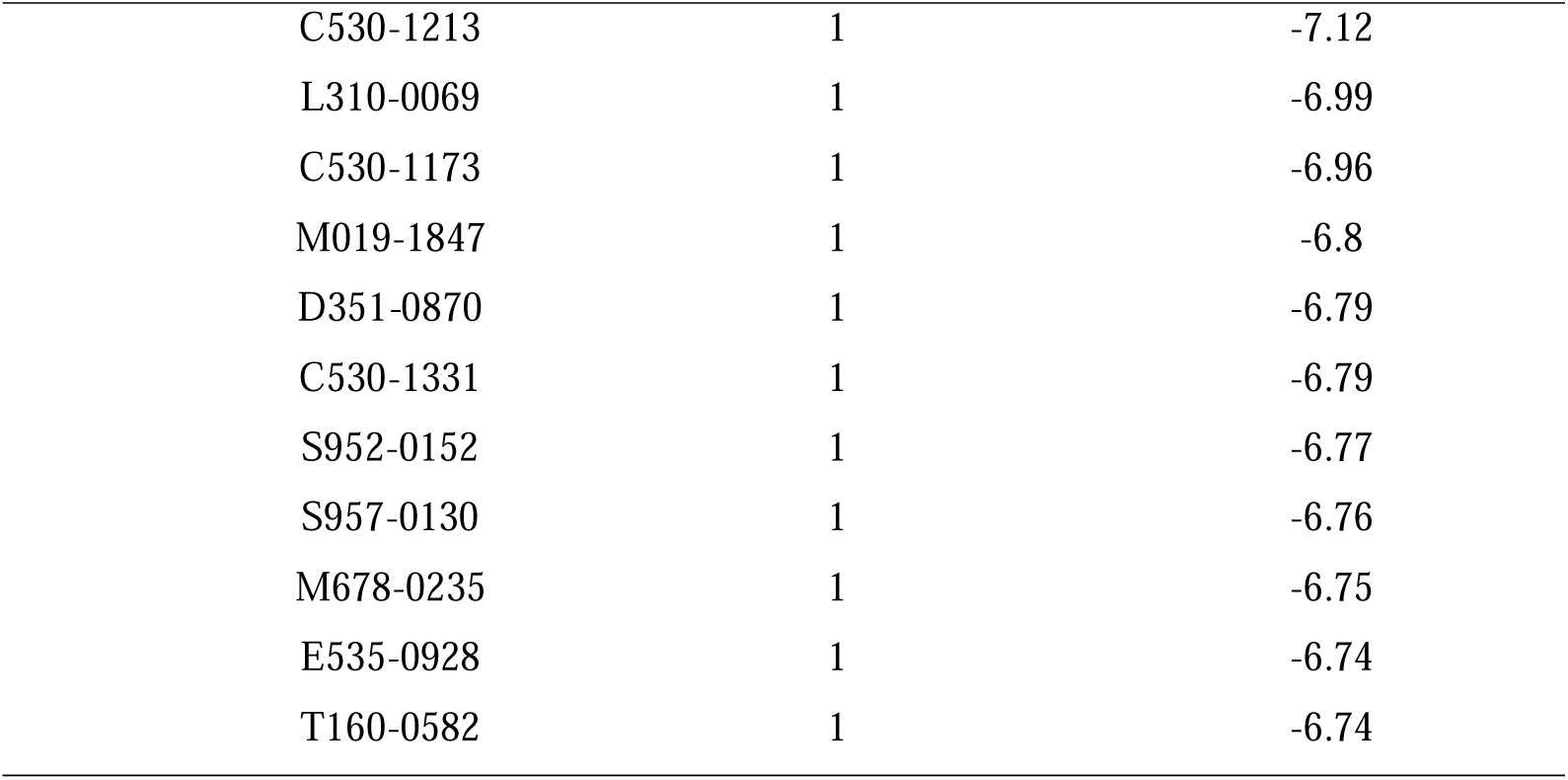
The representative screening result for target O14626, O95800, Q9HC97 by DeepGPCR and Schrödinger.

| Target | Name | DeepGPCR score | Schrödinger score<br>(Kcal/mol) |
| --- | --- | --- | --- |
| O14626 | 8015-6811 | 1 | -8.02 |
|  | T842-2224 | 1 | -6.97 |
|  | SC41-0196 | 1 | -6.9 |
|  | V007-8842 | 1 | -6.83 |
|  | S828-3346 | 1 | -6.6 |
|  | D399-0391 | 1 | -6.42 |
|  | SA92-0434 | 1 | -6.33 |
|  | J004-1117 | 1 | -6.19 |
|  | SC41-0276 | 1 | -6.13 |
|  | P094-1462 | 1 | -6.04 |
|  | SA70-0617 | 1 | -6 |
|  | F892-0669 | 1 | -6 |
|  | D399-0532 | 1 | -5.94 |
|  | E565-0464 | 1 | -5.92 |
|  | S827-4083 | 1 | -5.89 |
|  | M621-0356 | 1 | -5.88 |
| O95800 | 8019-4552 | 1 | -5.69 |
|  | 8020-4415 | 1 | -4.49 |
|  | 0708-0003 | 1 | -4.33 |
|  | 8011-6477 | 1 | -3.13 |
|  | Y031-6963 | 1 | -2.77 |
|  | 8020-6929 | 1 | -2.45 |
|  | 0896-4678 | 1 | -0.53 |
|  | 3137-0411 | 1 | -0.16 |
| Q9HC97 | K788-9238 | 1 | -8.07 |
|  | V030-8466 | 1 | -7.58 |
|  | K784-5585 | 1 | -7.47 |
|  | C529-0941 | 1 | -7.44 |
|  | S947-5093 | 1 | -7.41 |
|  | D233-0341 | 1 | -7.2 |
|  | S953-0095 | 1 | -7.16 |
|  | V004-3123 | 1 | -7.12 |

|  |  |  |
| --- | --- | --- |
| C530-1213 | 1 | -7.12 |
| L310-0069 | 1 | -6.99 |
| C530-1173 | 1 | -6.96 |
| M019-1847 | 1 | -6.8 |
| D351-0870 | 1 | -6.79 |
| C530-1331 | 1 | -6.79 |
| S952-0152 | 1 | -6.77 |
| S957-0130 | 1 | -6.76 |
| M678-0235 | 1 | -6.75 |
| E535-0928 | 1 | -6.74 |
| T160-0582 | 1 | -6.74 |

**Table S13.** The RMSD, TM-score between Alphafold2 predicted structure (Protein 1) and experimental PDB structure (Protein 2) for 62 selected GPCR. Here, we only selected sequence identity (ID)>=0.83.

| Protein 1<br>(F1-model_v4) | Protein 2<br>(From PDB) | Aligned<br>length | RMSD | TM-score | ID |
| --- | --- | --- | --- | --- | --- |
| AF-Q14416 | Q14416_4XAQ | 443 | 0.87 | 0.99 | 0.99 |
| AF-P41146 | P41146_4EA3 | 278 | 0.91 | 0.98 | 1.00 |
| AF-Q9Y271 | Q9Y271_6RZ4 | 290 | 1.23 | 0.72 | 0.99 |
| AF-P43220 | P43220_3C5T | 104 | 1.24 | 0.94 | 0.98 |
| AF-P21453 | P21453_3V2W | 295 | 1.27 | 0.65 | 0.92 |
| AF-P25103 | P25103_6E59 | 292 | 1.31 | 0.59 | 0.92 |
| AF-Q9UBS5 | Q9UBS5_4MQF | 406 | 1.34 | 0.97 | 1.00 |
| AF-P51686 | P51686_5LWE | 266 | 1.38 | 0.93 | 0.92 |
| AF-P41145 | P41145_6VI4 | 283 | 1.40 | 0.97 | 0.98 |
| AF-P47900 | P47900_4XNW | 297 | 1.50 | 0.83 | 0.98 |
| AF-P35408 | P35408_5YHL | 280 | 1.54 | 0.96 | 0.98 |
| AF-P51681 | P51681_4MBS | 295 | 1.57 | 0.82 | 0.95 |
| AF-P34972 | P34972_5ZTY | 301 | 1.60 | 0.65 | 0.94 |
| AF-Q92633 | Q92633_4Z34 | 307 | 1.62 | 0.77 | 0.93 |
| AF-Q9Y5Y4 | Q9Y5Y4_6D26 | 322 | 1.70 | 0.70 | 0.98 |
| AF-P31422 | P31422_2E4Y | 517 | 1.76 | 0.96 | 0.98 |
| AF-P30542 | P30542_5N2S | 284 | 1.78 | 0.72 | 0.95 |
| AF-P56726 | P56726_6O3C | 483 | 1.78 | 0.96 | 0.95 |
| AF-P29274 | P29274_2YDO | 295 | 1.79 | 0.90 | 0.98 |
| AF-P61073 | P61073_3OE9 | 270 | 1.81 | 0.62 | 0.94 |
| AF-P32300 | P32300_4EJ4 | 281 | 1.86 | 0.62 | 0.99 |
| AF-Q99835 | Q99835_4N4W | 357 | 1.87 | 0.76 | 0.97 |
| AF-Q92847 | Q92847_6KO5 | 293 | 1.88 | 0.70 | 0.97 |
| AF-P41594 | P41594_4OO9 | 244 | 1.88 | 0.58 | 0.94 |
| AF-P20789 | P20789_4XEE | 311 | 1.90 | 0.65 | 0.95 |
| AF-P25929 | P25929_5ZBQ | 285 | 1.92 | 0.59 | 0.93 |
| AF-P55085 | P55085_5NDD | 316 | 1.93 | 0.54 | 0.91 |
| AF-O43614 | O43614_4S0V | 319 | 1.96 | 0.64 | 0.88 |
| AF-P07550 | P07550_3D4S | 300 | 2.00 | 0.66 | 0.92 |
| AF-O00222 | O00222_6BT5 | 440 | 2.04 | 0.94 | 0.99 |
| AF-P28223 | P28223_6WHA | 244 | 2.05 | 0.91 | 0.96 |
| AF-P50052 | P50052_5UNF | 290 | 2.06 | 0.72 | 0.96 |
| AF-P32248 | P32248_6QZH | 276 | 2.07 | 0.36 | 0.93 |
| AF-P28222 | P28222_4IAQ | 309 | 2.17 | 0.80 | 0.87 |
| AF-P48039 | P48039_6ME2 | 297 | 2.17 | 0.59 | 0.87 |
| AF-P43116 | P43116_7CX2 | 265 | 2.19 | 0.91 | 0.96 |
| AF-P21730 | P21730_6C1Q | 299 | 2.19 | 0.76 | 0.95 |
| AF-P49286 | P49286_6ME6 | 308 | 2.19 | 0.66 | 0.89 |
| AF-P42866 | P42866_4DKL | 287 | 2.37 | 0.61 | 0.97 |
| AF-P51436 | P51436_6IQL | 275 | 2.37 | 0.75 | 0.86 |
| AF-P25090 | P25090_6LW5 | 302 | 2.38 | 0.67 | 0.97 |
| AF-P23385 | P23385_1ISS | 452 | 2.42 | 0.92 | 1.00 |
| AF-P30556 | P30556_4YAY | 285 | 2.45 | 0.67 | 0.93 |
| AF-P24530 | P24530_5XPR | 310 | 2.49 | 0.70 | 0.96 |
| AF-P35400 | P35400_2E4Z | 434 | 2.50 | 0.91 | 0.95 |
| AF-P32238 | P32238_7F8U | 314 | 2.52 | 0.67 | 0.88 |
| AF-Q9H244 | Q9H244_4NTJ | 272 | 2.65 | 0.67 | 0.89 |
| AF-P21917 | P21917_5WIV | 290 | 2.70 | 0.71 | 0.91 |
| AF-P08912 | P08912_6OL9 | 320 | 2.71 | 0.72 | 0.88 |
| AF-Q14832 | Q14832_3SM9 | 427 | 2.73 | 0.88 | 0.84 |
| AF-P30968 | P30968_7BR3 | 278 | 2.74 | 0.54 | 0.94 |
| AF-P41595 | P41595_5TUD | 322 | 2.79 | 0.78 | 0.87 |
| AF-P41143 | P41143_6PT3 | 301 | 2.84 | 0.69 | 0.93 |
| AF-P28335 | P28335_6BQG | 315 | 2.88 | 0.76 | 0.89 |
| AF-Q13255 | Q13255_3KS9 | 435 | 2.95 | 0.87 | 0.84 |
| AF-P08172 | P08172_3UON | 302 | 3.07 | 0.64 | 0.91 |
| AF-P08588 | P08588_7BTS | 307 | 3.21 | 0.62 | 0.90 |
| AF-P08483 | P08483_4U14 | 319 | 3.55 | 0.65 | 0.84 |
| AF-P14416 | P14416_6CM4 | 306 | 3.64 | 0.64 | 0.83 |
| AF-P35367 | P35367_3RZE | 311 | 3.70 | 0.65 | 0.85 |
| AF-Q14833 | Q14833_7E9H | 766 | 3.82 | 0.87 | 0.99 |
| AF-P47871 | P47871_5XEZ | 307 | 4.38 | 0.47 | 0.83 |
| Average |  | 322.24 | 2.19 | 0.74 | 0.93 |

## Notes

### Competing Interest Statement

The authors have declared no competing interest.

