## Supplemental materials for "Revolutionizing GPCR-Ligand Predictions: DeepGPCR with experimental Validation for High-Precision Drug Discovery"

**Supplementary material section 1**

**Schrödinger glide docking procedure**

We used Schrödinger glide docking to further screening compounds based on the predicted the binding affinity. First, the ligands underwent geometric optimization using the Ligprep module. The ligand energy was minimized using the OPLS 2005 force field, and all its ionized states were generated at pH 7.4. Subsequently, a single, low-energy three-dimensional structure of the ligand was generated, while preserving its original chiral state. Hydrogen atoms were added to the protein, and the system was optimized at pH 7.4 using the OPLS-u-2005 force field. The receptor grid was generated based on the geometric center coordinates of the protein’s ligands, with a size of 26 Å x 26 Å x 26 Å. The docking process adopted default settings such as standard precision, flexible sampling, and no constraints.

**Supplementary material section 2**

**Screening against three targets,** **O14626, O95800, and Q9HC97**

Overall, we selected three GPCR proteins, namely O14626 (GPR171), O95800 (GPR75), and Q9HC97 (GPR35), to showcase the applications of our DeepGPCR model in screening potential therapeutic compounds. These three GPCR proteins are all potential therapeutic targets for various diseases, including cancer, metabolic syndrome, and related disorders. For instance, GPR171 is a T-cell checkpoint that plays a crucial role in tumor immunity, inhibiting it can be a potential anticancer drug ^1^. Similarly, targeting the 20-HETE/GPR75 pathway has been discovered as a new, highly druggable potential target in the metabolic syndrome ^2^ and is also promising in interfering with prostate tumor cell malignant progression ^3^. Lastly, GPR35 is a potential target for various diseases ^4^, including cancer ^5^. Using our DeepGPCR model, we screened 102,472, 102,563, and 102,592 candidates with a score of ≥ 0.999 for O14626, O95800, and Q9HC97, respectively. We then grouped each protein's candidates into 1000 clusters and obtained potential representative compounds. Further clustering these 1000 compounds into 30 clusters helped us identify the final representative compounds depicted in Figure S6. The clustering analysis employed the default Ward's Hierarchical Agglomerative Clustering Method ^6,7^ implemented in Clusfps (https://github.com/kaiwang0112006/clusfps).

Considering the DeepGPCR model cannot directly provide GPCR-ligand binding conformation, we relied on the Schrödinger to dock the candidates into the protein targets. For O14626 and Q9HC97, we directly docked the 1000 representative candidates to their predicted pockets. However, the predicted binding cavity of O95800 was much narrower due to structural constraints according to our visual observation of the pocket region of its 3D structure. We tested the docking of molecules of different sizes. We observed that larger molecules had worse docking scores, which is consistent with our observation that the binding cavity is too narrow for large molecules to bind. Therefore, we selected 1923 compounds with a molecular weight ≤300 from its 102,563 candidate compounds for docking.

**Supplementary material section 3**

**Detailed procedure of pocket MD and metadynamics simulation**

The initial protein-compound complexes were from the top score conformation Schrödinger docking. The ligand was edited by PyMOL software ^8^ to make it in the correct protonation state at pH 7.

To make the simulation closer to GPCR environment, we have carried MD for GPR35-compound complexes embedded in lipid membrane. We preparation of the simulation system by building the GPR35-membrane-water-small molecule complex and solvate it using an appropriate solvent model. The system should be energy minimized and equilibrated using molecular dynamics simulation before the production run. Metadynamics simulations can estimate binding free energy calculation to explore whether protein-ligand will bind in solution. Metadynamics relies on adding a bias potential to sample the free energy landscape along a specific collective variable of interest ^9^,^10^. Note that the binding free energy calculations from Metadynamics may only be suitable for detecting the general trend of binding in virtual screening.

The MD simulation was carried out by Gromacs with AMBER-99 force field ^11,12^. The lipid PDB structure file (DPPC_293K.pdb)) and parameter files (DPPC.itp) are downloaded from Slipids websever (<http://www.fos.su.se/~sasha/SLipids/Downloads.html>) ^13–15^. The topology of the ligand and the partial charges of the ligand were generated by ACPYPE ^16^, which relies on Antechamber ^17^. The procedure to prepare the initial protein-ligand-lipid membrane with solvent and counter ion are largely follow the Gromac tutorial (http://www.mdtutorials.com/gmx/membrane_protein/02_topology.html), the major difference is that the forcefield we used was AMBER instead of GROMOS in the tutorial. We used TIP3P water molecules ^18^, and the counter ions were added to neutralize the total charge using the Gromacs program tool ^19^. The long-range electrostatic interactions under the periodic boundary conditions were calculated with the Particle Mesh Ewald approach ^20^. A cutoff of 10 Å was used for van der Waals non-bonded interactions. Covalent bonds involving hydrogen atoms were constrained by applying the LINCS algorithm ^21^.

We performed the energy minimization steps with a step-size of 0.001ns, 100 ps simulation with an isothermal-isovolumetric ensemble (NVT), and 10ns simulation with the isothermal-isobaric ensemble (NPT) for water equilibrium. After that, a 40ns NPT production run (step size 2 fs) was carried out. The Parrinello-Rahman barostat and the modified Berendsen thermostat were used for simulation with a fixed temperature of 308 K and a pressure of 1 atm. RMSD and hydrogen bond number of the trajectory were calculated using Gromacs tools.

The simulation continued using the metadynamics approach to explore the free energy landscape. We carried 40ns metadynamics simulation with Plumed^22^ patched Gromacs. The protein-ligand complex's interface coordination number of atoms was used as a collective variable (CV). The protein-ligand interface coordination numbers correlate with the numbers of atom contact, and a larger coordination number usually indicates that the protein-ligand is binding.

The coordination number C is defined as follows by Plumed:

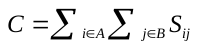
 (1) and

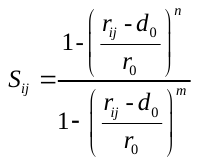
 (2)

In the simulation, n was 8, m was 12, $d_{0}$ was 0 nm, and $r_{0}$ was 0.25 nm. $d_{0}$ is a parameter of the switching function. $r_{ij}$ is the distance between atom i and atom j. The degrees of contact between two groups of atoms can be estimated by the above function(1) ^22^. Metadynamics simulation for each protein-ligand system was performed for 40 ns. During the metadynamics simulation, Gaussian values were deposited every 1 ps with a height of 0.3 kJ/mol. The widths of the Gaussians were 5 for the coordination number. The free energy landscapes of the metadynamics simulations along the CV were generated by the Plumed program and plotted using Gnuplot ^23^.
